# Rapamycin mitigates Valproic Acid-induced teratogenicity in human and animal models by suppressing AP-1-mediated senescence

**DOI:** 10.1101/2023.08.29.555421

**Authors:** Giovanni Pietrogrande, Mohammed R. Shaker, Sarah J. Stednitz, Farhad Soheilmoghaddam, Julio Aguado, Sean Morrison, Samuel Zambrano, Tahmina Tabassum, Ibrahim Javed, Justin Cooper-White, Thomas P. Davis, Terence J O’Brien, Ethan K. Scott, Ernst J. Wolvetang

## Abstract

Valproic acid (VPA) is an effective and widely used anti-seizure medication but is teratogenic when used during pregnancy, affecting brain and spinal cord development for reasons that remain largely unclear. Here we designed a genetic recombinase-based *SOX10* reporter system in human pluripotent stem cells that enables tracking and lineage tracing of Neural Crest cells (NCCs) in a human organoid model of the developing neural tube. We found that VPA induces extensive cellular senescence and promotes mesenchymal differentiation of human NCCs at the expense of neural lineages. We next show that the clinically-approved drug, Rapamycin, inhibits AP1-mediated senescence and restores aberrant NCC differentiation trajectory in human organoids exposed to VPA. Notably, *in vivo* validation in developing zebrafish highlighted the therapeutic promise of this approach. Collectively our data identifies a novel mechanism for VPA-associated neurodevelopmental teratogenicity and a potential pharmacological preventative strategy. The results exemplify the power of genetically modified human stem cell-derived organoid models for drug discovery and safety testing.

## Introduction

Valproic acid (VPA) is widely prescribed for the treatment of bipolar disorder, migraine, and epilepsy [1]. It is the most effective anti-seizure medication (ASM) for the treatment of generalised epilepsy syndromes [2, 3] and influences voltage-dependent sodium channels, GABA neurotransmitter release, HDAC-mediated epigenetic modifications, modulation of neurotropic factor production, and dendritic spines re-organization [4]. However, its use during pregnancy leads to an increased risk of teratogenesis termed Fetal Valproate Syndrome (FVS, OMIM #609442). FVS encompasses heart defects, severe craniofacial defects, neural tube defects, and neurocognitive disorders. Consequently, VPA is considered contraindicated for use during pregnancy in many countries [5], and prescription to women of childbearing potential is not recommended even in absence of plans for pregnancy. Regrettably, numerous patients lack alternative therapeutic options with equivalent therapeutic effectiveness [6, 7]. Studies in zebrafish, mice, rats, chickens, and human cellular model have shown similar VPA teratogenic effects [8–12], suggesting a largely evolutionary conserved fundamental pathogenic process. Strikingly, Neural Crest cells (NCCs) are involved in the development of most embryonic tissues affected by VPA.

During development NCCs migrate from the neural tube and differentiate into diverse cell types, including melanocytes, peripheral neurons, and smooth muscle cells [13]. This developmental plasticity extends to mesenchymal stem cells (MSCs), further emphasizing their potential in contributing to various tissue formations, but remarkably few reports have investigated the impact of VPA exposure on NCC. VPA alters avian NCC proliferation and migratory potential *in vitro* [14], and in zebrafish affects the development of the palate, a structure derived from the progeny of NCCs [15]. Nevertheless, the cellular and molecular processes associated with these events remain unclear.

To elucidate the mechanism of VPA-induced teratogenicity, here we engineered a *SOX10:mMaple-fLPo/eGFP* human induced pluripotent stem cell (hiPSC) line that incorporates a genetic switch to facilitate detection and lineage tracing of SOX10^+^ NCC across multiple lineages. This approach allows us to detect variations in physiologically relevant behaviour of NCC in 3D organoid models of human neural tube derived spinal cord (hSCOs). Our findings demonstrate that VPA exposure promotes inappropriate differentiation of newly specified NCCs and point to a strong connection with the observed widespread rise in cellular senescence. Indeed, the senomorphic drug Rapamycin [16] demonstrates substantial efficacy in preventing this increase in senescence and associated defects in NCC differentiation. Remarkably, *in vivo* zebrafish model replicates the VPA-induced NCC differentiation events and the subsequent Rapamycin rescue, providing additional validation for the biological significance of our findings beyond the *in vitro* setting. Finally, we identify the master transcription factor AP1 as a likely mediator of VPA-induced senescence and aberrant differentiation, rather than VPA activity as an epigenetic modulator. All together these results shed light on the delicate balance between cellular senescence and cell fate choices during human VPA teratogenicity, proposing a potential pharmacological prevention strategy through Rapamycin treatment.

## Results

### Conditional genetic switch for *SOX10* gene expression report and lineage tracing

We designed a genetic switch driven by *SOX10*expression that upon activation leads to the expression of a *SOX10*-driven reporter and constitutive expression of a second reporter for permanent lineage tracing of the progeny derived from SOX10+ NCCs. The first element is directly dependent on *SOX10* transcription, and leads to the concomitant expression of a fluorescent reporter (mMaple [17]) and the DNA recombinase Flp [18]. We used Cas9-mediated gene targeting to edit the 3’ UTR region of *SOX10*, 125 bp after the stop codon to insert an IRES-mMaple-P2A-fLPo expression cassette (Fig1a and SFig1a). The second element (Fig1b) was randomly integrated into the genome and drives constitutive expression of a Neomycin selectable module that, after excision by Flp [19], enables constitutive GFP-expression based lineage tracing. We verified the functionality of this switch by differentiating the targeted bulk hiPSC population into Schwann Cells Precursors (SCPs), a *SOX10* expressing cell type. As expected mMaple expression was functionally confirmed by photoconversion (SFig1c).

**Fig1.**
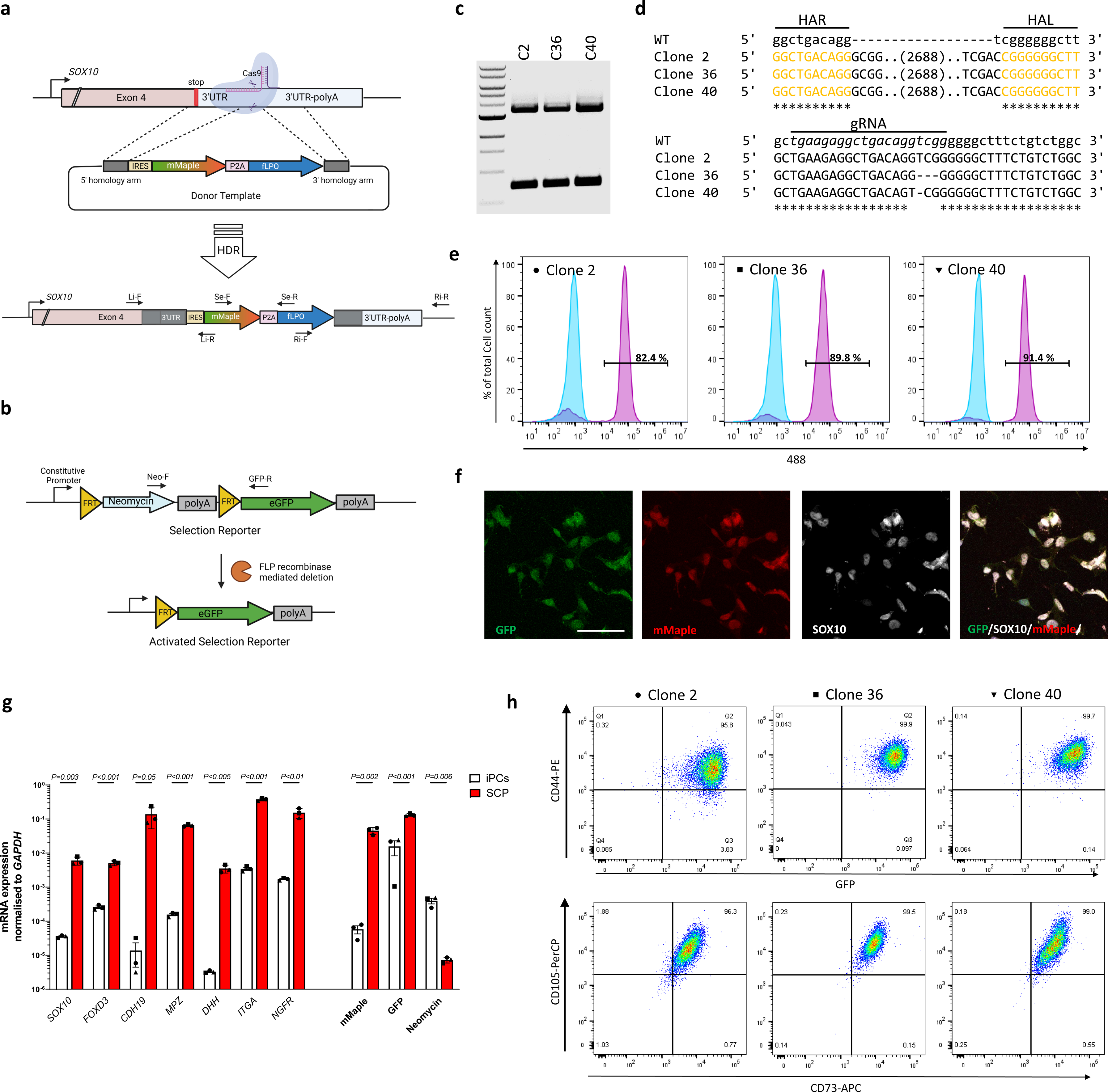
Generation of specific and lasting genetic switch driven by *SOX10*. **a** Schematic representation of the 3’ UTR region of the *SOX10* locus. gRNA cutting site is indicated by scissors. Plasmid donor is designed to insert in the *SOX10* 3’ UTR region a reporter (mMaple)/recombinase (FLPo) cassette with an IRES sequence. Indicated primer pairs are used in the following characterizations. **b** Schematic of the selection reporter cassette that is activated by *SOX10*-driven FLPo recombinase expression. **c** Three clones were randomly selected and screened for switch insertion within the locus using the primers F/Ri-R. The higher band shows bp weight expected by monoallelic integration, and the lower band shows bp weight expected by the intact locus. 1Kb ladder. **d** Upper band from C was purified and sequenced on both the left (Li-F/Li-R) and right (Ri-F/Ri-R) sides to verify homologous recombination. Lower band from C was purified to verify the presence of indels. gRNA sequence in italic. **e** Emission in the 488 channel assessed by FACS of each iPC clone without (blue histogram) and after (purple histogram) differentiation protocol. **f** Representative image of the immunostaining for SOX10 on SCPs at the end of the differentiation protocol. Scale bar 10 µm. **g** Relative RNA abundance in differentiated SCPs of the indicated SCPs markers and the indicated genetic switch element normalized to *GAPDH* mRNA and compared to the respective initial iPC clone. Each dot represents one clone. Error bars represent s.e.m.; two-tailed Student’s t test. **h** SCPs derived from each clone were subsequently further differentiated into Mesenchymal stem cells. At the end of the differentiation protocol cells were tested by FACS for emission in the 488 channel and expression of a panel of mature MSCs markers to verify their cellular identity. Isotypic control incubation on primary human MSCs was used to set up negative gating.

### *SOX10* genetic switch is activated upon differentiation into *SOX10* expressing cells, is tightly regulated and preserved across multiple cell types

Starting from the bulk population of edited iPSCs containing the genetic switch (SFig1b and c) we isolated Clone 2, Clone 36, and Clone 40 with correct integration of the *mMaple/fLPo* cassette (Fig1c and d). Clone 36 and Clone 40 presented indels on the other *SOX10* allele, generated during the Cas9 gene editing process (Fig1d). We again functionally evaluated the efficiency of the switch in each clone by differentiation to SCPs. The genetic switch was specifically activated in SOX10 positive cells only (Fig1e and f and SFig1d). Importantly, neither the original iPSC bulk population nor any of the iPSC clones presented a detectable amount of 488 fluorescence positive cells (Fig1e, blue histogram and SFig1b), highlighting the strong specificity of our system that is only activated by substantial expression of endogenous *SOX10*. Robust differentiation is shown by the sharp decrease of pluripotency markers (SFig1e) and upregulation of a panel of SCPs markers. This is accompanied by changes in expression of the genetic switch elements (Fig1g), that are indicative of the Flp recombinase-driven excision of the Neomycin resistance transgene. This result was further supported by the selective death of GFP positive SCPs induced in the presence of G418 (SFig1f). To validate the robustness of lineage tracing across multiple lineages we differentiated SCPs from each of the three clones, sorted the GFP positive population and further differentiated the resulting population into MSCs. We found that more than 99% cells were positive for MSC markers, lost mMaple expression and retained GFP expression for at least 12 weeks and after further differentiation [8] (Fig1h and SFig2).

### Neural crest cells originate and differentiate in a 3D model of neural tube development

3D organoid models can recapitulate developmental signalling cascades more closely than a 2D system, and are an excellent platform to study human development, drug toxicity or teratogenicity [20]. We adapted a recently described protocol to generate neural tube derived spinal cord organoid (SCO) [21] from human iPCS (Fig2A). Wholemount immunostaining for BRN2 showed the establishment of an elongating neuroepithelial layer after neuromesodermal induction and following 4 days of bFGF (SFig4a). We next observed the appearance of GFP/mMaple positive cells (Fig2b and 2c) after only 6 days of bFGF, culminating at day 12 (SFig4b). Wholemount immunostaining confirmed that GFP/mMaple positive cells express SOX10 (Fig2d). Interruption of FGF, and exposure to retinoic acid (RA) for 4 days promoted loss of SOX10/mMaple expression (Fig2e) and SOX10+ cells differentiation towards ISL1/2^+^ neurons (Fig2f, g and h), mirroring events of *in vivo* ventral neural tube development [22], as previously also reported [21]. We confirmed our observations in three independent organoid batches; qPCR analysis shows increased expression of multiple trunk NCC markers [23] (Fig2i) and genetic switch elements (Fig 2l) over time, that are turned off after retinoic acid (RA) addition in favour of markers of neuronal lineage. Next, we leveraged mMaple and GFP expression to study the behaviour of NCC within SCOs in real time (Video1). Over 6 days we observed both *de novo* specification (SFig3a) and proliferation (SFig3b) of mMaple+ NCCs. Using AI-based segmentation [24] and tracking software [25] we tracked GFP positive cells and identified two groups of mMaple positive and negative cells that locate in different regions (SFig3c and SFig3d). Interestingly, mMaple positive cells move with higher speed (SFig3e), as expected for highly migratory cells. We conclude that our lineage tracing approach together with live imaging enables high throughput and scalable screening and allows the identification of compounds that can perturb the behaviour and/or cell identity of cells of interest within human organoid models.

**Fig2.**
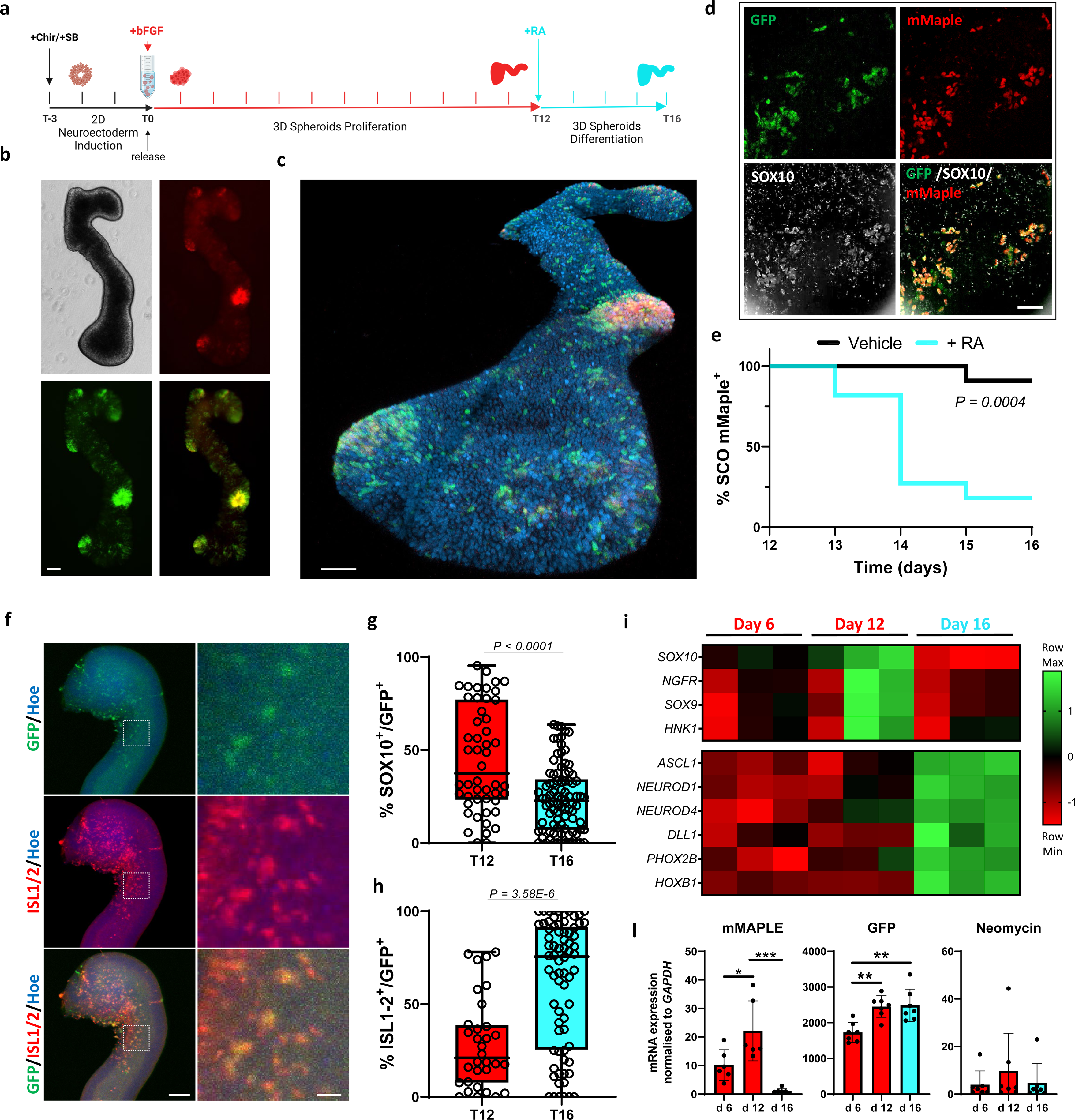
SOX10 driven genetic switch is activated in SCOs and enables lineage tracing. **a** Schematic representation of SCO differentiation protocol. Stages are divided into 2D/neuroectodermal induction, 3D proliferation/specification and 3D differentiation. **b** Representative live image at day 10 taken with tabletop fluorescent microscope Ti2U (Nikon) of an organoid where cells with activated genetic switches are indicated by emission in the 488nm (green) and 555nm (red) spectra after photoconversion and overlay. **c** 3D reconstruction of the same organoid as **b** imaged at super-resolution after fixation. Scale bar 100 µm. **d** Representative field of whole mount of photoconverted d12 SCO immunolabelled for SOX10 (white) and overlay with eGFP (green) and mMaple (red). Scale bar 50 µm. **e** Kaplan–Meier curve of mMaple positive SCOs. Organoids were imaged every day from day 12 to 16 during exposure to retinoic acid (n=11) or vehicle (n=11) and evaluated for the presence of photoconverted (mMaple positive) cells within the organoid. Long-rank test. **f** Representative field of whole mount d16 SCO immunolabelled for ISL1/2 (red) and overlay with eGFP (green). Left scale bar 70 µm, right scale bar 35 µm. **g** Box plots show the percentage of SOX10 and **h** ISL1/2 positive cells on the total number of GFP positive cells. Each data point in the box plot represents an ROI of an organoid section where GFP cells were present. Whiskers represent min-max values; d12 n=33, d16 n=67 ROI were analysed per organoid; two-tailed Student’s t test. **i** Total RNA from 5-7 hSCOs pooled was extracted at different stages of differentiation and used to quantify the mRNA expression levels of the indicated neural crest cells and neuronal markers after normalized to *GAPDH* mRNA. The significance threshold was set to P=0.01 as determined by one-way ANOVA with Tukey’s multiple-comparison post-hoc corrections. Each column in the heatmap represents one of three independent differentiation batches. **l** Total RNA from 3-5 pooled hSCOs of 5 to 7 replicates was extracted at different stages of differentiation and used to quantify the mRNA expression levels of the indicated genetic switch element normalized to *GAPDH* mRNA. Error bars represent s.d.; two-tailed Student’s t test, * P<0.05 ** P<0.01, *** P<0.001.

### Valproic acid (VPA) causes premature differentiation and SASP (senescence associated secretory pathway) induced cellular senescence

After establishing that we developed a system that reliably reports on the presence and differentiation of NCCs, we investigated the aetiology of neural tube malformations caused by VPA during the proliferation phase (T0 to T12). After 72 hours of treatment with FGF and 2mM VPA (a dosage with clinical relevance [26], subsequently referred to as VPA unless otherwise specified), 82% of organoids did not show detectable mMaple signal (Fig3a and SFig4c). This decrease in SOX10:mMaple/fLPo expression was confirmed at transcriptional level (Fig3b) and was associated with a decrease in the expression of NCC markers (SFig4d) that was remarkably consistent across a range of VPA concentrations (Fig4e). Bulk RNA sequencing performed on SCOs generated from the three previously described clones C2, C36 and C40 treated with VPA revealed downregulation of a panel of NCC mRNAs (Fig3c). Intriguingly, *SOX10* was the second most significantly deregulated gene following VPA exposure (*Padj=*3.59E-39). Further analysis of the bulk RNA seq revealed a substantial upregulation of a large pool of senescence associated genes [27] (Fig3d) and enrichment for KEGG pathways closely associated with proliferation, autophagy and inflammation (Fig3e), like MAPK and AKT pathways, also supported by increases in phospho-ERK and phospho-AKT levels (SFig4f) [28]. To test the biological significance of this increase in senescence we next exposed SCOs generated from three different clones to VPA and VPA + Rapamycin, an efficient SASP inhibitor [29] (Fig3f). We found that Rapamycin prevented the VPA-induced upregulation of senescence marker p21 (Fig3g) and increased activity of senescence associated (SA)-β-Galactosidase (Fig3h). As expected, Rapamycin affected the expression of SASP [30] induced by VPA (Fig3i) and other processes like proliferation and DNA damage repair, hallmarks of cellular senescence (SFig4g). These results suggest that dysregulated SASP expression and anomalous senescence during development may contribute to the teratogenic properties of VPA and are prevented by Rapamycin.

**Fig3.**
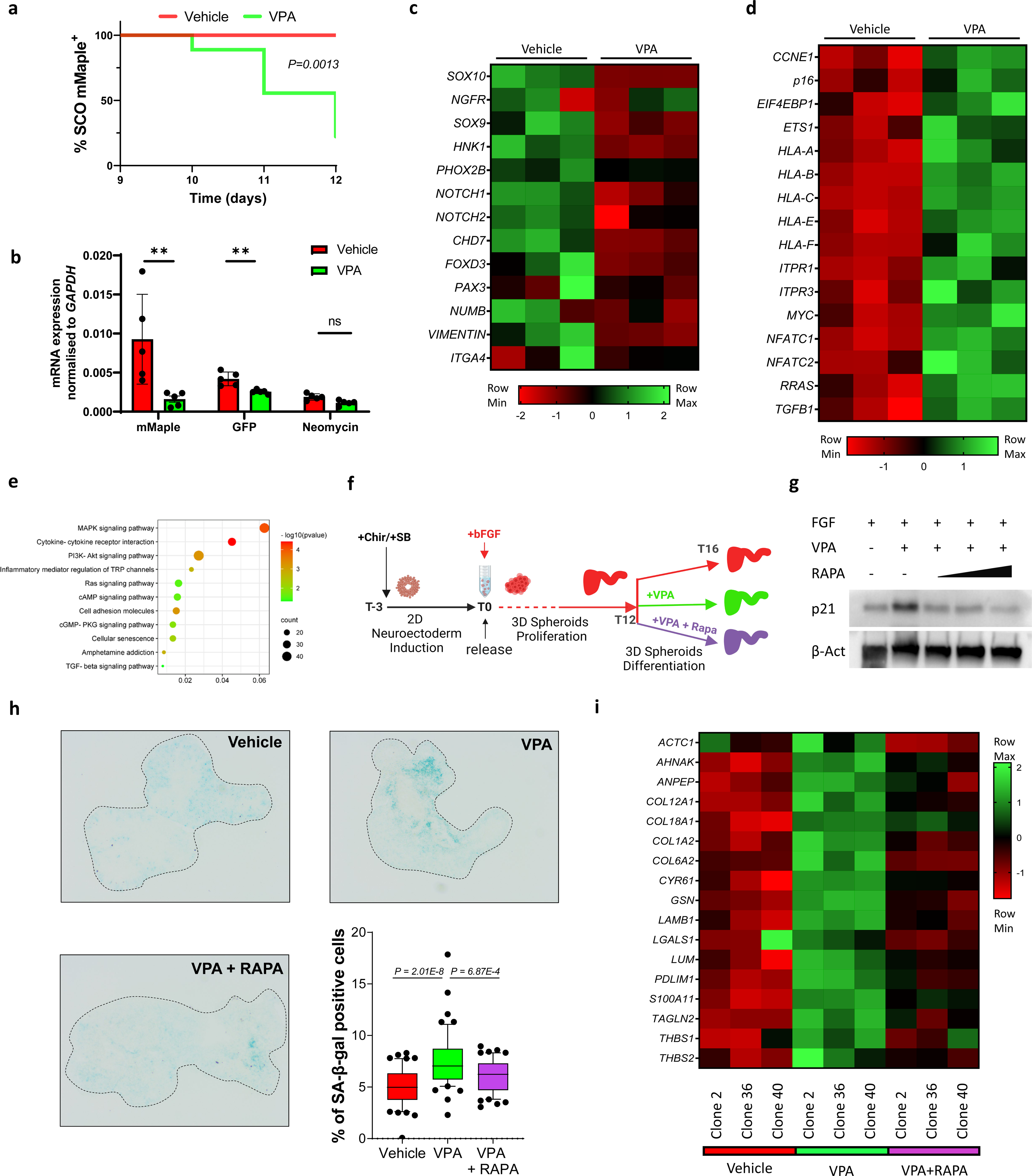
VPA induces NCC differentiation and promotes cellular senescence. **a** Kaplan–Meier curve of mMaple positive SCOs. Organoids were imaged every day from day 9 to 12 during exposure to vehicle (n=10) or VPA (n=9) and evaluated for the presence of photoconverted (mMaple positive) cells within the organoid. Long rank test. **b** Total RNA from 3-5 pooled hSCOs of 5 replicates exposed to vehicle or VPA was extracted at different stages of differentiation and used to quantify the mRNA expression levels of the indicated genetic switch element normalized to *GAPDH* mRNA. Error bars represent s.d.; two-tailed Student’s t test, ** P<0.01. **c** 5-7 hSCOs generated from three independent clones were pooled, total RNA was extracted after exposure to vehicle or VPA and used for bulk RNA sequencing to investigate the expression levels of the indicated neural crest cells markers and **d** senescence markers. Each column in the heatmap represents one of three clones. **e** Gene Set Enrichment Analysis was carried out using DAVID. The statistically significant senescence signatures were selected (FDR < 0.25) and placed in order of enrichment. Each pathway is enriched in VPA treatments as compared to vehicle-treated organoids. Heatmap was plotted by https://www.bioinformatics.com.cn/en, a free online platform for data analysis and visualization. **f** Schematic representation of SCOs experimental design. **g** Western blot analysis for organoids exposed to FGF alone (vehicle), VPA and VPA and increasing concentrations of Rapamycin, 125, 250 and 500 mM left to right. **h** SA-β-gal assays were performed on organoid sections. Each data point in the whisker box plot represents a single section analysed. Error bars represent s.d.; at least 12 individual organoids were analysed per condition; one-way ANOVA with Tukey’s multiple-comparison post-hoc corrections, the significance of each comparison is indicated in the graph. **i** Heatmap showing Senescence-Associated Secretory Phenotype (SASPs) expression generated from bulk RNA sequencing of organoids exposed to vehicle, VPA and VPA + Rapamycin identified on SASP atlas as per **c**.

**Fig4.**
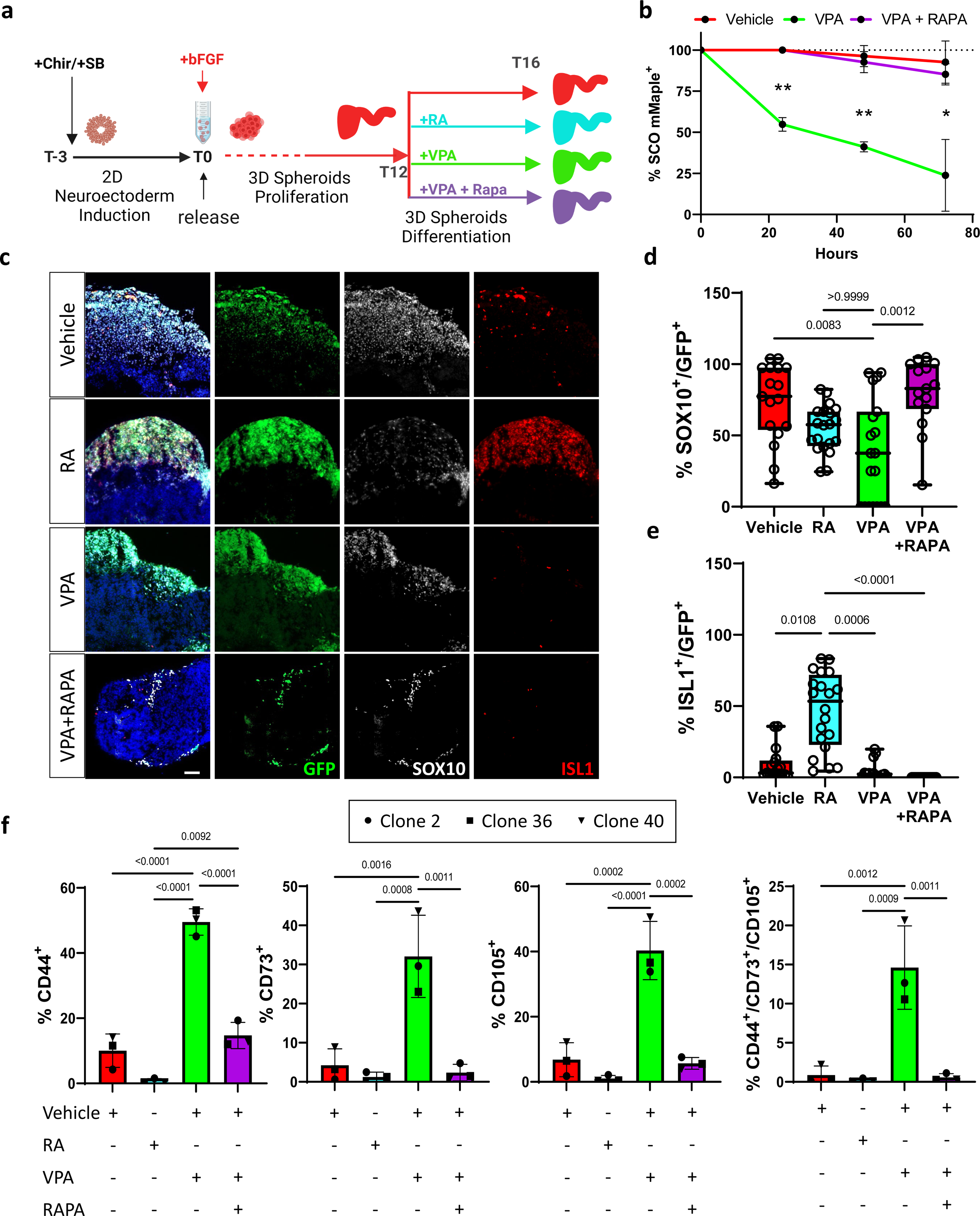
VPA promotes NCC differentiation into Mesenchymal Stem cells. **a** Schematic representation showing the timeline of organoids exposure to the indicated compounds and combinations. **b** Graph showing the percentage of mMaple positive SCOs over time during exposure to the indicated compounds. Organoids were imaged every day from day 12 to 16 during exposure to vehicle (C2 n=7, C36 n=6, C40 n=6), VPA (C2 n=7, C36 n=7, C40 n=8), or VPA+Rapamycin (C2 n=9, C36 n=9, C40 n=9), and evaluated for the presence of photoconverted (mMaple positive) cells within the organoid. Two-way ANOVA analysis attributed a significant variation to treatment (53,42%, p=0.0001) and time (20,75%, p=0.0004). The significance of simple effects within each timepoint is indicated in the graph. **c** Representative fields of sections of organoids after multiple treatments immunolabelled for SOX10 (white) and ISL1 (red). Scale bar 50 µm. **d** Quantification of the colocalization of SOX10 with GFP and **e** ISL1 with GFP presented in **c**. Each point of the box plot represents an ROI on a section. 8-12 organoids generated from each clone and pooled were analyzed; one-way ANOVA with multiple comparison post-hoc correction. **f** Quantification of FACS analysis for cells presenting Mesenchymal stem cells markers and quantification of triple positive % in organoids generated from each clone after exposure to multiple compounds. RAPA refers to VPA+Rapamycin. Error bars represent s.d.; One-way ANOVA with multiple comparison post-hoc correction.

### Rapamycin prevents precocious differentiation of NCCs towards mesenchymal lineage caused by VPA

After we identified senescence as a potential driver of NCCs differentiation, we aimed to dissect NCC fate choices, and test whether Rapamycin could prevent this process (Fig4a). We observed that VPA caused precocious NCC differentiation in SCOs generated from all three clonal lines, as indicated by the loss of mMaple positive cells. Importantly, this was prevented by Rapamycin (Fig4b) without significantly altering the total number of GFP+ cells (SFig4h). We next leveraged our lineage tracing to unravel the identify the progeny derived from NCCs (Fig4c). As expected, in vehicle SCOs most of GFP+ positive cells express SOX10 (Fig4d), while after exposure to RA these SOX10+ cells switch towards the neuronal lineage (Fig4e). VPA treatment substantially decreased the presence of SOX10+ cells but did not lead to an increased presence of neuronal cells (Fig4e and SFig4i). Indeed, genes significantly repressed (*Padj*<0.05) after VPA exposure are mostly expressed in neurons, as indicated by their mapping to the neuronal cluster (SFig4j) in the single cell transcriptomic atlas of human spinal cord [31]. Remarkably, Rapamycin preserved the SOX10+ population, and thus prevented the precocious differentiation of NCCs induced by VPA exposure. Transcriptional data from bulk (Sfig5a) and clonal iPSC line (SFig5b) derived organoids treated with VPA revealed an increased expression of MSC markers. Similarly, genes significantly induced by VPA exposure (*Padj*<0.05) map to a foetal mesenchymal progenitors’ cluster (SFig5c) identified by scRNA sequencing of human foetal spinal cord [32]. However, these gene expression changes were not observed after exposure to RA (SFig5d). Rather, VPA exposure impaired the specification of NCCs-derived neuronal cells (SFig5e). In support of these data, FACS analysis confirmed that, after VPA, GFP+ cells begin to express the canonical MSCs markers CD44, CD73 and CD105 (Fig4f and SFig6). Intriguingly, only VPA treated SCO presented a conspicuous number of GFP cells positive for all three markers, while Rapamycin prevented this differentiation. Importantly, GFP negative cells presented a negligible level (0.5%<) of triple positive population in all the conditions analysed (Sfig7). Thus, we conclude that VPA exposure causes endogenous NCCs to begin differentiating into MSCs. This notion is further corroborated by the observation that GFP+ cells derived from dissociated SCOs could attach, proliferate, and survive in MSC permissive medium only after exposure to VPA (SFig5e and f).

### Rapamycin prevents precocious differentiation of NCCs caused by VPA in developing zebrafish

To confirm that Rapamycin treatment can prevent VPA induced differentiation of NCCs in developing animals, we exposed fertilised zebrafish eggs to vehicle, VPA and VPA + Rapamycin from fertilization until hpf 48. We noticed that most of the controls broke through the embryo chorion while VPA treated embryos often failed to achieve this (results not shown). Defects in the development of embryos exposed to VPA were macroscopic and severely affected the development of the craniofacial area, dorsal orbit and the eye (Fig5a) [33]. Whole mount immunostaining for sox10 (Fig5b) revealed a significant decrease in sox10+ cells in all regions analysed (Fig5e), including the areas of palate (Fig5C) and neck/head (Fig5a). Interestingly, Rapamycin exposure was sufficient to prevent the loss of sox10+ cells induced by VPA. We further noticed evident changes in the morphology of NCC derived melanocytes [34] (Fig5f) that were also rescued by rapamycin treatment, further supporting our working hypothesis that VPA alters the behaviour of NCC and their progeny that are prevented by Rapamycin.

**Fig5.**
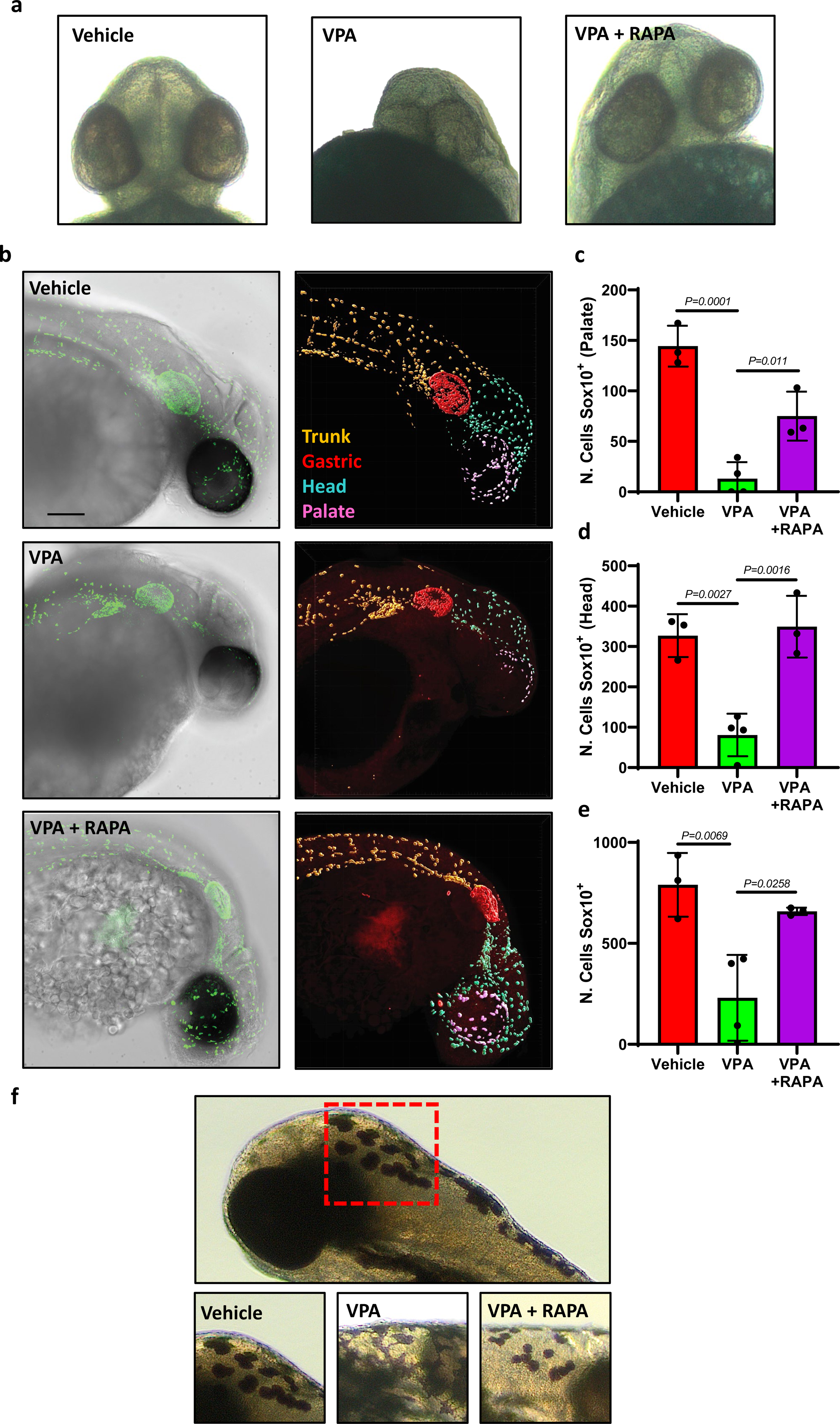
Rapamycin prevents VPA-induced promotion of NCC differentiation in Zebrafish. **a** Representative image of the cranial region of zebrafish larvae at 48 hpf exposed to the indicated compounds since 2 hpf. **b** Representative images of bright field overlayed with confocal imaging of zebrafish larvae exposed to multiple compounds and hybridised with an antibody for Sox10 (green) and 3D reconstruction of identified Sox10+ cells within the larvae. The upper body was partitioned into multiple regions, with each cell being assigned a specific color corresponding to its localized area. Scale bar 100 µm. **c** Quantification of the number of Sox10 positive cells counted in the palate region (purple), head (**d**) and entire upper body (**e**) of zebrafish larvae exposed to vehicle, VPA and VPA+Rapamycin. Each point of the bar graph represents one larvae. Error bars represent s.d.; one-way ANOVA with multiple comparison post-hoc correction. **f** Representative image of the morphology of melanocytes in the cervical area of larvae after exposure to the indicated compounds.

### AP1 mediates VPA induced defects and Rapamycin rescue

Finally, we pinpointed the molecular mechanism causing VPA associated differentiation of NCC. Starting from our bulk RNAseq analysis (SFig8a) we initially identified a substantial effect of Rapamycin on recovering the transcriptional profile induced by VPA exposure (SFig8b and c) and we shortlisted a pool of potential target genes overexpressed after VPA exposure but repressed by Rapamycin (*Padj*<0.05, Fig 6a) which pointed to the involvement of transcriptional regulators (Fig6b). We found that a number of pathways related to CNS development and senescence were altered (SFig8e). Among the most significant differentially expressed genes we identified a pool of pioneer factors carrying the potential to cause substantial transcriptional changes [35] that are restored to basal level by Rapamycin (Fig6c). Using STRING online tools [36] we discovered that mTOR and these pioneer factors are closely associated on multiple levels through FOS (Fig6d and SFig8d) and point to an involvement of the AP1 (Fos/Jun) master transcriptional factor. Importantly we verified that other proteins belonging to the Fos/Jun network follow a similar transcriptional pattern after exposure to VPA and Rapamycin (SFig8f). To confirm that AP1 is a mediator of VPA induced senescence, we exposed SCOs generated from a randomly chosen clone (Clone 40) to VPA and VPA + SR11302, a specific inhibitor of AP1 (iAP1) that does not alter the activity of VPA as histone acetylase inhibitor (SFig8g). We found that specific AP1 inhibition prevented the loss of mMaple+ organoids (Fig6e), caused a sharp decrease in sen-β-Galactosidase activity (Fig6f) and p21 positive cells (Fig6g) and prevented the loss of SOX10+ cells (Fig6h) similarly to Rapamycin. Altogether these findings suggest that AP1 is the key element that is driving the VPA induced senescence, and the inhibitory effect of Rapamycin is due to the repression of AP1.

**Fig6.**
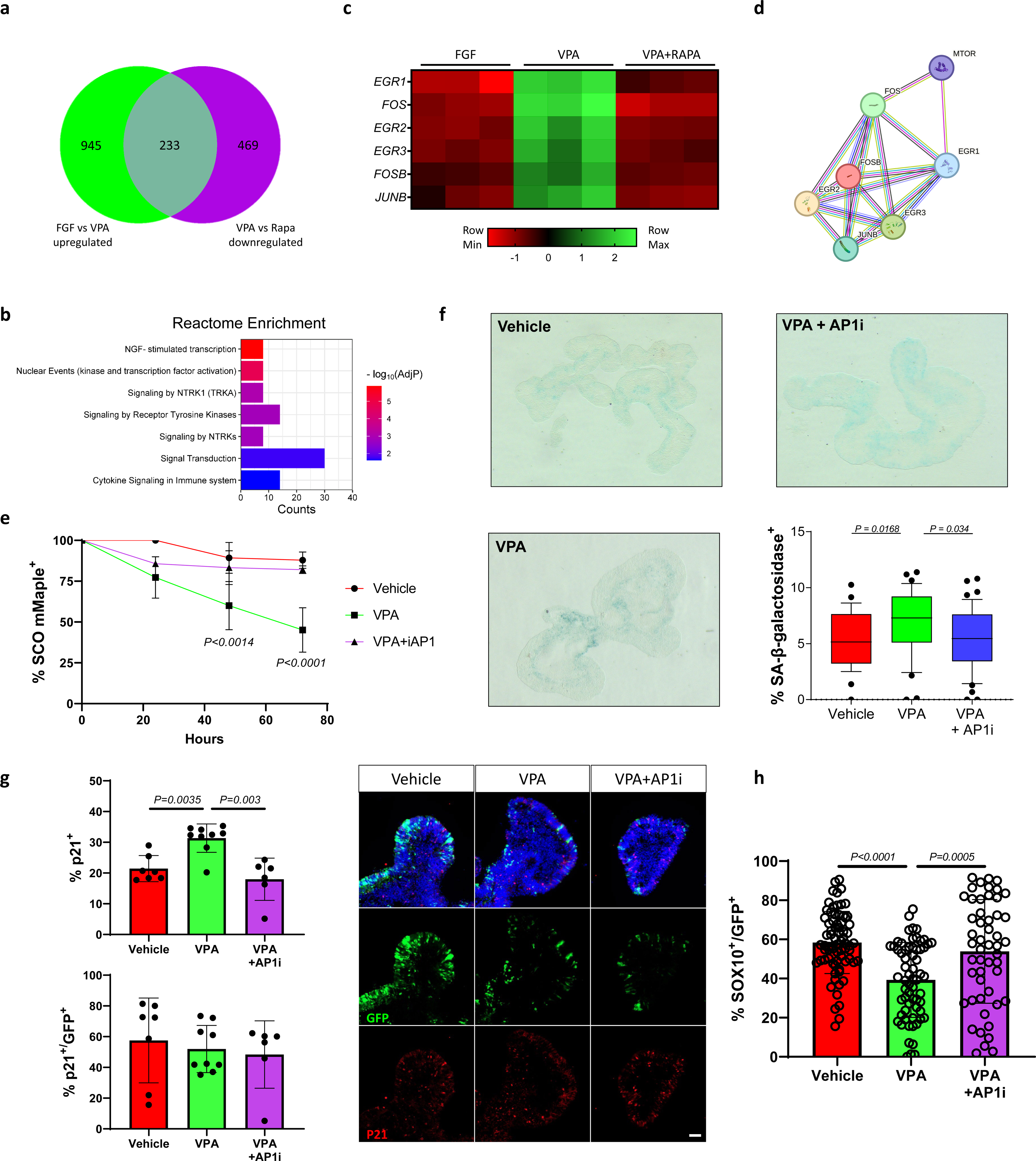
VPA-induced senescence is dependent on AP-1 activity. **a** Ven diagram shows the gene population significantly upregulated upon VPA exposure (Green, Vehicle vs VPA) but differentially repressed by Rapamycin (Purple, VPA vs Rapamycin). **b** Reactome enrichment analysis was carried out on the overlapping gene group identified in **a**. Bars indicate the REACTOME pathways identified, counts show the number of genes. **c** heatmap expression of the top 7 significantly differentially expressed genes of the 83 previously identified. **d** Interactome map of the shortlisted genes identified and relationship to mTOR. Known interactions bars: Blue, from curated database; Pink, experimentally determined; Yellow, textmining; Black, co-explression; Purple, protein homology. **e** Graph showing the percentage of mMaple positive SCOs over time during exposure to the indicated compounds. Organoids were imaged every day and two to 5 fields of view showing 4 to 12 organoids per condition per day were evaluated for the presence of photoconverted (mMaple positive) cells within each organoid. Two-way ANOVA analysis attributed a significant variation to time and treatment (P<0.0001). Multiple comparison significance of VPA group to other groups per each timepoint is shown on the graph. **f** SA-β-galactosidase assay was performed on organoid sections. Each data point in the whisker box plot represents the percentage of β-galactosidase positive cells in a single section analysed. 7-9 dividual organoids were analysed per condition. **g** Representative fields of sections of organoids after multiple treatments immunolabelled for p21 and quantification. **h** Quantification of SOX10/GFP positive cells in hSCOs after exposure to multiple treatments; 25-33 individual organoids were analysed per condition one-way ANOVA with Tukey’s multiple-comparison post-hoc corrections, the significance of each comparison is indicated in the graph. Error bars represent s.d.

## Discussion

In this study we show for the first time that senescence can be a driver of VPA-induced teratogenicity in developing human spinal cord. Importantly we also demonstrate *in vitro* and *in vivo* that VPA exposure severely affects the normal differentiation trajectory of neural crest cells, a cell population of pivotal importance during neurodevelopment, and this process can be prevented treatment with the senomorphic drug, Rapamycin.

Epilepsy is diagnosed in >5 million people each year, and VPA is one of the most widely adopted and most effective anti-seizure medications, in particular for people with generalised epilepsies [2, 3]. Despite the teratogenic effects of VPA being a major limitation to its use in clinical practice in women of childbearing potential with epilepsy, very little is known of the mechanisms underlying this. Features of VPA teratogenicity include neural tube defects, neurocognitive and neurobehavioural deficits, congenital heart defects, peculiar facial features, urogenital malformations and musculoskeletal abnormalities [37]. In this study we decided to focus our attention on Neural Crest Cells (NCCs), which arise during the first stages of neural tube development and contribute to the development of most tissues affected in FVS.

In order to dissect the mechanisms for the teratogenic effects of VPA in NCCs, we designed a genetic switch for gene expression reporting/lineage tracing. This switch is turned on by endogenous *SOX10* expression, does not display detectable leakage, and is maintained after multiple passages and across multiple lineages, thus overcoming the main limitations to lineage tracing in human cultured iPSC associated with oscillations in the expression of driver genes and inaccurate tracing [38]. This system enabled for the first time real time single cell tracking of the behaviour of spontaneously formed NCC in a 3D model of human developing neural tube [39]. Our approach has multiple advantages: it simplifies the detection of newly specified NCCs in 3D, identifies differentiation events as indicated by loss of the mMaple reporter, and allows lineage tracing of the differentiated progeny. Using this tool we confirmed in a human 3D system that RA exposure induces the differentiation of neural crest cells into ISL1+ peripheral neurons. Indeed, animal studies in chick and mouse suggest that NCCs differentiation is driven by a gradient of Retinoic Acid in vivo [22, 40, 41], and our results indicate that our model can successfully recapitulate this process, and thus is suitable to investigate VPA teratogenicity and NCC behaviour.

We observed that VPA significantly promoted the NCCs within the organoid to differentiate, switching from a proliferative to a differentiation program. This was accompanied by a significant induction of cellular senescence as shown by increased presence of multiple senescence markers including p16, MYC and TGFβ, SA-β-Galactosidase activity and p21 protein. Together our data are in strong agreement with the evolving body of literature supporting the pivotal role of cellular senescence in the precise regulation of developmental processes [42] and are in line with a previous study showing that neuroepithelial cells in mice and human organoids exposed to VPA display increased cellular senescence [43]. VPA is a histone deacetylase (HDAC) inhibitor, however the role of HDACs in senescence is still controversial, as there are reports of their involvement in both promoting [44] or hindering senescence [45]. We believe this is most likely explained by the specificity of single HDACs to different epigenetic targets. Regardless, our results suggest that VPA-induced senescence can be countered despite the HDAC-inhibition activity. Putting together the pieces of the epigenetic puzzle behind senescence is a next fundamental step to understand the phenomenon of aging.

Since HDAC inhibition has been previously associated with increased SASP (Senescence-Associated Secretory Phenotype) expression [46], and to establish a causal relationship between the induction of senescence and altered NCCs differentiation, we aimed to prevent VPA-induced senescence by targeting SASP [47]. We chose Rapamycin, an inhibitor of the mTOR signalling cascade and a senomorphic drug that has shown consistent effects on cellular senescence and substantial efficacy in different aging models [48]. Strikingly Rapamycin treatment resulted in a significant reduction in VPA-induced abundance of p21, a primary mediator of developmental senescence [49], and a drastic decrease in NCCs differentiation. In rodents premature senescence within the neural tube is associated with developmental defects that are prevented by Rapamycin [50]. Thus, we decided to confirm our findings in an *in vivo* model of developing animal.

Exposure of zebrafish larvae to VPA is not generally associated with neural tube defects, but other neurodevelopmental defects such as deformation of eyes and craniofacial abnormalities [51]. Intriguingly, NCCs contribute to the development of these tissues. Therefore, these observations support out view of a pivotal role of NCC in mediating the teratogenicity of VPA. Our results show that VPA treatment severely alters anatomical facial features with a concomitant and significant decrease in Sox10 positive cells. Similar defects are observed after ablation of HDAC4 [52]. Strikingly Rapamycin could largely prevent these defects, fully supporting our *in vitro* observations.

Finally, we investigated the molecular cascade leading to NCCs differentiation after VPA exposure. Transcriptome comparisons of VPA and Rapamycin treated organoids suggested an involvement of the AP1 network. AP1 is a master transcription factor involved in a wide range of processes, from growth to differentiations, and importantly AP1 is a pioneer factor that coordinates the cellular senescence program at a transcriptional level [53, 54]. Our results indicate that VPA-induced senescence is not NCCs specific, which could explain the diverse range of organ systems affected by VPA teratogenicity, and that AP1 inhibition can prevent both the increase in cellular senescence and the differentiation of NCC associated with VPA, without affecting its role as a HDAC inhibitor. Thus, AP1 is a key mediator of VPA induced senescence and neurodevelopmental teratogenicity. Interestingly, retinoids are repressors of AP1 activity [55] and given the presence of RA in an anteroposterior gradient along the developing spinal cord, our results suggest that dose-dependent VPA-induced defects affect most commonly the caudal region (i.e. spina bifida) as consequence of the lower concentration of RA in the area – which is supported by clinical studies [26, 56].

To our knowledge this is the first report linking VPA exposure to a specific alteration in the neural crest program of differentiation. One of the main characteristics of FVS is a distinctive facial appearance, which suggests a disruption of the normal differentiation and migration program of NCCs contributing to the craniofacial bone. However past literature has mainly focused on the effects of VPA on brain development, and indeed VPA exposure is a well-established method to generate rodent model of autism and neurocognitive deficits [57]. Importantly, our results suggest a dysregulated senescence program behind the teratogenic effects of VPA and identify the master transcription factor AP1 as the driver of the observed defects. Collectively, we propose an entirely novel model of VPA’s mechanism of teratogenicity, and suggest that co-treatment with Rapamycin could allow safe prescription of VPA to women of childbearing potential in whom it is the most effective medication, in particular those with generalised epilepsies.

## Materials and methods

### Plasmids

We inserted a reporter construct into WTC iPCs to express a mMaple-P2A-fLPo cassette [17] in the 3’ untranslated region of *SOX10*. pSpCas9(BB)-2A-Puro (PX459) V2.0 was a gift from Feng Zhang (Addgene plasmid # 62988) was modified to target the region of interest. The fLPo recombinase target used for lineage tracing purposes was a gift from Prof Ryan Lister [19].

### Cell culture and reagents

The WTC iPCs cell line was maintained in mTesr plus (100-1130, Stemcell Tech) after coating with Matrigel (354277, Stemcell Tech). Plasmidic constructs were delivered by nucleofection (P5 Primary Cell 4D, 197187 Lonza). Edited cells were enriched by a temporary incubation with 0.5 ug/ml Puromycin (Gibco). Single clones were generated by seeding single cells in presence of CloneR (05888, Stem Cell Tech). All clones and the bulk population were routinely subjected to mycoplasma testing. Schwann cell precursors (SCPs) were generated by exposing cells to a cocktail of SB-431542 10 μM (1614, Tocris), CHIR99021 3 μM (4423, Tocris), holo-transferrin 10 μg/ml (T0665 Sigma), heregulin 10ng/ml (78071, Stemcell Tech) Ascorbic acid (50 μg/ml), bFGF 8ng/ml (233-FB, R&D Systems) in DMEM F12 (11320033, Thermo Fisher), 2% B27 (Gibco, 17504-044), 1% Glutamax (35050061, Thermo Fisher Scientific) 1% MEM-NEAA (Gibco, 11140-050), 0.2% 2-Mercaptoethanol (Gibco, 21985), 1% penicillin and streptomycin (Life Tech), for 3 weeks. SCPs were plated on Matrigel coated multiwell at 10000cell/cm^2^ density and induced to differentiate into MSCs by changing the medium composition to low glucose DMEM (Thermo Fisher, 11885084), 10% FBS (Gibco), 1% penicillin and streptomycin for three weeks. Expression of MSCs markers was confirmed using the FACS Human MSC Analysis Kit (AB_2869404, BD Bioscience) according to the manufacturer instruction. Cells were analysed with a BD LSRFortessa followed by FlowJo analysis. The resulting MSCs were further differentiated to osteoblast and adipocytes using complete osteogenic media: DMEM low glucose (11885084, Gibco), 10% foetal bovine serum (Gibco), 1% penicillin-streptomycin, 100 nM Dexamethasone (265005, Merk) + 100 µM Ascorbic acid 2-phosphate (699004, Sigma) + 10 mM β-glycerolphosphate (G6501, Sigma), and complete adipogenic medium: α-MEM (A1049001, Thermo Fisher Scientific) supplemented with 0.5 mm isobutylmethylxantine (IBMX, I5879, Sigma)), 1 μm dexamethasone, 10 μm insulin (91077C, Sigma) and 200 μm indomethacin (405268, Merk), respectively.

Spinal cord organoids were generated as per [21] with minor modifications. Small iPSC colonies were plated and after 24 hours exposed to N2 mesodermal differentiation media 1% N2 (17502001, Gibco), 2% B27, 1% MEM NEAA, 0.1% 2-Mercaptoethanol, 1% penicillin and streptomycin, supplemented with SB-431542 10 μM (1614, Tocris) and CHIR99021 3 μM (4423, Tocris). After 3 days, colonies were detached by dispase incubation and grown in suspension in N2 differentiation media. The medium was supplemented daily with bFGF 20ng/ml (233-FB, R&D Systems), RA 1µg/ml (R2625, Sigma), VPA 1mM (OOAU00071, Sapphire bioscience), Rapamycin 200nM (US1553210, Merk), iAP1 SR11302 1µM (2476, Tocris) unless otherwise specified.

### Organoid sectioning and histology

Spinal cord organoids were fixed in 4% paraformaldehyde (PFA) for 1 hour at 4°C and washed with phosphate-buffered saline (PBS) three times for 10 minutes each at RT before allowing to sink in 30% sucrose at 4°C overnight and then embedded in OCT (Agar Scientific, cat. #AGR1180) and cryosectioned at 14 μm with a Thermo Scientific NX70 Cryostat. Tissue sections were used for immunofluorescence and for the SA-β-Gal assay. For immunofluorescence, sections were blocked and permeabilized in 0.1% Triton X-100 and 3% Bovine Serum Albumin (BSA) in PBS followed by incubation in a humid chamber with primary antibodies overnight at 4°C, washed three times with PBS 0.1% Triton X-100 and incubated with secondary antibodies for 1 hour at RT. 10 ng/ml DAPI (Sigma, cat. #D9564) was added for 5 minutes to mark nuclei. Secondary antibodies conjugated to Alexafluor 488, 568, or 647 (Invitrogen) were used for detection. For wholemount staining organoids were permeabilized by overnight incubation in PBS 0.5% Triton X-100 at 4°C, followed by the same immunostaining protocol.

### Zebrafish wholemount

Zebrafish were maintained in standard housing conditions as described in [58]. Tupfel long-fin wild-type embryos were collected at 0 hours post fertilization (hpf) and exposed to standard embryo medium (control) 150 µM VPA or 150 µM VPA and 1 mM Rapamycin for 48 hours. Embryos were then rinsed in embryo medium, manually dechorionated, and fixed in 4% PFA at 4°C overnight. Samples were incubated with primary antibody (anti-Sox10, 1:200) overnight at 4°C, washed, and incubated with secondary antibody (Alexa 488, 1:500) for 4 hours at RT. Immediately prior to imaging, embryos were mounted on slides in 2% low melting temperature agarose.

### Imaging and analysis

Immunofluorescence images were acquired using a Zeiss LSM 900 Fast Airyscan 2 super-resolution microscope, Zeiss AxioScan Z1 Fluorescent Imager, PerkinElmer Operetta CLS High Content Analysis System, and Yokogawa W1 Spinning Disk Confocal for zebrafish embryos. Images were analysed with a custom pipeline compiled on CellProfiler or ImageJ. Cells in each frame of the time-lapse were segmented using the AI-based software CellPose [24] using the cytosolic GFP channel signal. The position (centroid) of the segmented cells in each frame were used for cell tracking by applying the Hungarian linker algorithm through custom-made MATLAB software, as described in [25].

### Molecular analysis

For western blot analysis samples were lysed in RIPA buffer (Thermo Fisher) followed by one sonication pulse (1’’ at 40% intensity, VCX750, Sonics). The protein content was assessed by BCA assay (23225, Thermo Fisher). 50 to 100 ug protein was loaded in each lane and separated on 4–20% precast polyacrylamide gel (17000436, Biorad) and transferred on PVDF membrane (1704156, Biorad) using Transblot Turbo (Biorad) program 1. The membrane was blocked with 5% skim milk in TBS-T for 1 hour and incubated with primary antibodies overnight at 4°C, washed and incubated with secondary antibodies for 1 hour at RT. After brief exposure to ECL substrate (1705060, Biorad) the luminescent signal was detected with a ChemiDoc MP Imaging system (Biorad).

### Antibodies

SOX10 (89356, NEB, 1:400), sox10 zebrafish (AB229331, ABCam, 1:200), BRN2, p21 (2946, NEB, 1:3000), Isl1/2 (67.4E12, DSHB, 1:50), OP (ab8448, ABCam 1:1000), FABP4 (ab92501, ABCam 1:500), B-Actin (A3864, Sigma, 1:10000), ERK (9102, NEB, 1:10000); phospo-ERK (9101, NEB, 1:10000); AKT (9272, NEB, 1:5000); Phospo-AKT (9271, NEB, 1:5000); cFOS (134122, ABCam, 1:2000), ; anti-rabbit IgG (Invitrogen, A10042, 1:500); anti-rabbit IgG (Invitrogen, A21245, 1:500); anti-mouse IgG (Invitrogen, A11029, 1:500); anti-mouse IgG (Invitrogen, A21235, 1:500).

### RNA isolation

RNA from cells or spinal cord organoids was extracted with Nucleospin RNA (Scientificx) according to the manufacturer’s instructions. RNA integrity was evaluated with the 2100 Bioanalyzer RNA 6000

Pico Chip kit (Agilent) using the RNA Integrity Number (RIN). RNA samples with a RIN > 7 were considered of high enough quality for bulk RNA sequencing.

### Bulk RNA sequencing

Messenger RNA was purified from total RNA using poly-T oligo-attached magnetic beads. After fragmentation, the first strand cDNA was synthesized using random hexamer primers. Differential expression analysis between conditions was performed using the DESeq2 R package (1.20.0). The resulting P-values were adjusted using the Benjamini and Hochberg’s approach for controlling the false discovery rate. Genes with a *Padj* <=0.05 found by DESeq2 were assigned as differentially expressed. Gene Ontology (GO) enrichment analysis of differentially expressed genes was implemented by the cluster Profiler R package or using online tools [59].

### Real-time quantitative PCR

1 to 0.5 μg of total RNA was reverse transcribed using iScript cDNA Synthesis Kit (Bio-Rad). A volume corresponding to 5 ng of initial RNA was employed for each real-time PCR reaction using PowerUp SYBR Green Master Mix (Applied Biosystems) on a CFX Opus Real-Time PCR detection system. Glyceraldehyde-3-phosphate dehydrogenase (GAPDH) was used as normaliser. Primers sequences (5’-3’ orientation) are listed in STable 1.

### Statistical analysis

Results are shown as mean ± standard deviation (s.d.) or Box Plot as median and whisker. P value was calculated by the indicated statistical tests on Prism software. In the figure legends, n indicates the number of independent experiments or biological replicates when not specified in results.

## Competing interests

The authors declare no competing interests.

## Acknowledgments

We thank Novogene for performing bulk RNA sequencing and associated bioinformatic analysis; the analytical facility and the histology facility at the School of Miomedical Sciences, University of Queensland for the technical support, Dr Rob Sullivan at the Queensland Brain Institute Histology Core for advice on zebrafish whole-mount immunohistochemistry, the Queensland Brain Institute Advanced Microscopy Facility, Prof Ryan Lister at the University of Western Australia for providing the BLADE constructs, Dr Alison Anderson and Dr Huiwen Zheng for comments and suggestions on the transcriptomic data. This project was in part funded by NHMRC-MRFF-Stem Cell Mission APP 2007653, SJS and EKS were supported by a Simons Foundation Research Award (625793), TJO is supported by an NHMRC Investigator Grant (APP1176426). JA was supported by the Swiss IBSA Foundation for scientific research, the Jérôme Lejeune Foundation and a NHMRC Ideas Grant (2001408), MRS was supported by the Children Hospital Foundation (PCC0252021), GP was supported by European Leukodystrophy Foundation International (ELA-2021-024F2).

## Contributions

GP, MS and SM generated human spinal cord organoids. GP, MS, SS, FS, JA, SM, TT, JCW, TD, ES and EW contributed to acquisition, analysis or interpretation of data. SS and IJ generated the *in vivo* data. SZ analysed in vivo live tracing data. GP and EW contributed to experimental design, planned and supervised the project and wrote the paper. All authors edited and approved the final version of this article.

## Supplementary figure legends

**Sfig1**

**a** PCR primers were designed to map on the 5’ and 3’ ends of the switch integrated in the *SOX10* locus and sequenced to verify homology driven repair and incurrence of potential indels. **b** FACS sorting measuring the activation of the genetic switch expressing GFP over the differentiation protocol from iPCs to SCPs. **c** Representative live image showing colocalization of GFP (green) and photoconverted mMaple (red) in SCPs. **d** Quantification of GFP, mMaple and SOX10 positive SCPs generated from the three indicated clones and rate of colocalization of relevant markers. **e** Quantification by qPCR of iPCs markers for pluripotency *OCT4*, *NANOG* and *SOX2* in each clone at the iPCs stage (white) and after differentiation into SCPs (red). Each symbol represents a different clone; two-tailed Student’s t-test. error bars represent s.d. **f** Representative field of live fluorescent microscopy of SCPs after 6 days incubation with G418.

**Sfig2**

**a** Schematic representation of the differentiation protocol of iPCs into SCPs and further differentiation into MSCs, adipocytes and osteoblasts. **b** Representative live image of MSCs generated from bulk iPCs derived SCPs after 7 days of differentiation protocol. The channels refer to the excitation frequency of GFP (488nm) and mMaple (562nm). The image shows emission before and after photoconversion. **c** Histogram FACS analysis to identify GFP+ population of MSC generated from each clone (red peak) compared to the original iPCs population (azure peak). The relative percentage of GFP+ cells is indicated in each graph. After 3 weeks, more than 90% of MSCs were GFP positive, consistent with the GFP percentage detected at the initial SCP stage. **d** qPCR quantification of SCPs markers expression *SOX10* and *FOXD3* in independent MSCs differentiation of SCPs generated from three clones. **e** Expression levels of MSCs markers *CD44*, *CD90* and *CD105* after differentiation. **f** Expression levels of transgenes associated with the genetic switch mMaple, GFP and Neo measured by qPCR in SCPs and MSCs from three independent clones. **g** Representative image of MSCs (CTRL) differentiated into osteoblasts (OSTEO) and processed for immunofluorescent staining for Osteopontin (OSTP). Graphs are showing relative % of GFP positive cells untreated (CTRL) and after differentiation for all three clones and the quantification of Osteopontin expression in each independent differentiation. Quantifications refer to pulled values of three replicates per clone. **h** Representative bright filed image of MSCs and osteoblasts stained with Azalin red and quantification of the staining by absorbance in three replicates for each clone. **i** Representative images of MSCs and MSC-derived Adipocytes immunolabelled for Fatty Acid-Binding Protein 4 (FABP4, red) and quantification of GFP positive cells and FABP4 in untreated (CTRL) and after differentiation for all three clones. Quantifications refer to pulled values of three replicates per clone. Notably, the undifferentiated MSCs population retained strong GFP expression after 3 weeks from specification. **l** Representative bright field image of MSCs and Adipocytes stained with Oil red and quantification of the staining by absorbance in three replicates for each clone. Significance is indicated per each column; Two-tailed Student’s t-test, error bars represent s.d.

**SFig6**

**a** Region of interest of whole organoid live imaging at the beginning and at the end of the session of 96 hours. Photoconversion identifies two groups of GFP positive cells, one mMaple positive and the other mMaple negative, and was renewed every 24 hours to identify newly specified neural crest cells. **b** Single cells were tracked over time. **c** Quantification of displacement over time of every tracked cell (upper panel). Gradient color shows the degree of displacement on the indicated axis for GFP (middle) and mMaple (lower panel) cells. **d.** Left panel, green indicates the trajectories of GFP only cells, while red indicates GFP/mMaple positive cells. The right panel shows the movements of each tracked cell identified with a different color. **e** Box plot quantification of speed and ratio of displacement/time for mMaple positive and negative cells (in a.u.), Two-tailed Student’s t-test.

**SFig4**

**a** Representative immunostaining for neuroepithelial marker Brn2 on elongating neural tube during proliferation phase. **b** Quantification of the percentage of organoids presenting detectable photoconvertible mMaple signal. Analysis of 3 replicates n= 8-15 organoids. **c** Representative live imaging of hSCO over the last 4 days of the proliferation phase, control and exposure to Valproic acid. Green shows emission at 520nm, red shows emission at 600nm after UV-induced photoconversion. Bar 50 µm. **d** Quantification by qPCR for the presence of transcripts of the NCC markers *SOX10*, *SOX9* and *HNK1* in samples extracted from 3 independent batches of hSCOs generated from bulk iPCs population, n=3-6 HSCOs were pulled for each sample. Two-tailed Student’s t-test, error bars represent s.d. **e** Western blot analysis for SOX10 protein content in HSCOs exposed to increasing concentrations of Valproic acid. β-ACTIN was used as protein loading control. 8-12 organoids were pulled for each lysate and 100 µg were loaded per lane. **f** Western blot analysis for the indicated proteins and phosphorylated forms in hSCOs exposed to increasing concentrations of Valproic acid as per **d**. **g** GO term enrichment analysis showing the Top 8 biological processes emerging from the list of genes differently expressed between VPA and VPA+Rapamycin groups. **h** HSCO were desegregated by incubation with Accutase, resulting cells were passed through a mesh strain to remove doublets and analysed by FACS analysis. Plot shows an example of gating used to differentiated GFP negative and positive cells. Bar graph shows the percentage of GFP positive cells present in hSCOs exposed to the indicated compounds, in three replicates each with n=4-7 individual organoids pooled. One-way ANOVA, error bars represent s.d. **i** Relative quantification by qPCR for the presence of the indicated markers of neuronal lineage in hSCOs exposed to VPA. n=3-6 hSCOs were pulled for each sample; two-tailed Student’s t-test, error bars represent s.d. **j** Bulk RNAseq identified genes significantly downregulated by VPA (*Padj*<0.05) mapping on single cell spinal cord transcriptomic database. Gene set mean expression shown in color scale.

**SFig5**

**a** Relative RNA expression in 3-5 pooled hSCOs of 5 replicates exposed to vehicle or VPA of MSCs markers normalized to *GAPDH* mRNA. Error bars represent s.d.; two-tailed Student’s t test, ** P<0.01. **b** Heat map shows selected MSC markers measured as transcriptomic expression across three hSCOs independent replicates exposed to vehicle or Valproic Acid. **c** Bulk RNAseq identified genes significantly upregulated by VPA (*Padj*<0.05) mapping on single cell transcriptomic of 13^th^ week post-fertilization stage human male spinal cord, donor 8. Fetal mesenchymal progenitor cluster is circled in red. Gene set mean expression shown in color scale. **d** Total RNA was extracted from 3 groups of 4-5 pooled hSCOs generated from bulk iPCs at different timepoints during the differentiation and the expression of the indicated MSCs markers was examined by qPCR and quantified as relative to *GAPDH*; One-way ANOVA, error bars represent s.d. **e** hSCOs were exposed to 4 days of RA after 12 days of FGF or 8 days FGF/4days FGF+VPA, then sectioned and stained for the neuronal marker ISL1. White arrows show loss of colocalization with GFP. **f** Schematic representation of the experimental plan showing the timeline of exposure to different compounds in hSCO medium (N2 medium plain) or MSCs medium, without any coating, and exemplificative field image at bright field and at 488nm excitation of cells from disaggregated hSCOs after the indicated time in culture. The red cross indicates that GFP-expressing cells did not attach without coating. GFP-positive cells could attach and proliferate if the surface was coated with Matrigel (not shown). **g** FACS analysis for the indicated MSC markers of GFP positive cells derived from hSCOs exposed to VPA, disaggregated and grown in culture in MSC medium for three weeks.

**SFig6**

Individual FACS analysis of each clone and each marker used for the quantification shown in fig 4f.

**SFig7**

Individual FACS analysis for the expression of MSCs markers of the GFP negative fraction of fig 4f for each clone, and quantification. Gating was kept equal for GFP positive and negative analysis.

**SFig8**

**a** Venn diagram shows differentially expressed transcripts between hSCOs exposed to vehicle (FGF), VPA and VPA+Rapamycin (RAPAMYCIN) with a significance *AdjP* value <0.05. **b** Heatmap and unsupervised clustering of the whole transcriptome by bulk RNA sequencing of the indicated samples. **c** Principal component analysis of the three replicates of each condition used during bulk RNA sequencing and subsequent analysis. **d** Western blot analysis of protein samples extracted from hSCOs exposed to the indicated compounds for the expression of cFOS and loading control shown by β- ACTIN. **e** GO term enrichment analysis of differentially expressed genes identified as significantly differently expressed between Vehicle and VPA groups (left) and between VPA and VPA+Rapamycin groups (right). **f** Network of genes closely associated by interaction at multiple levels to FOS and genes belonging to the FOS network showing significantly differential expression after the indicated treatments. The number of readings was extracted from the bulk RNA sequencing data. **g** Representative Western blot analysis for the content of acetylated H3 (Ac-H3), histone H3 and β- ACTIN in protein samples extracted from hSCOs exposed to the indicated compounds.

## Notes

### Competing Interest Statement

The authors have declared no competing interest.

