## Supplementary figures and images for "Rapamycin mitigates Valproic Acid-induced teratogenicity in human and animal models by suppressing AP-1-mediated senescence"

### Supplementary figures

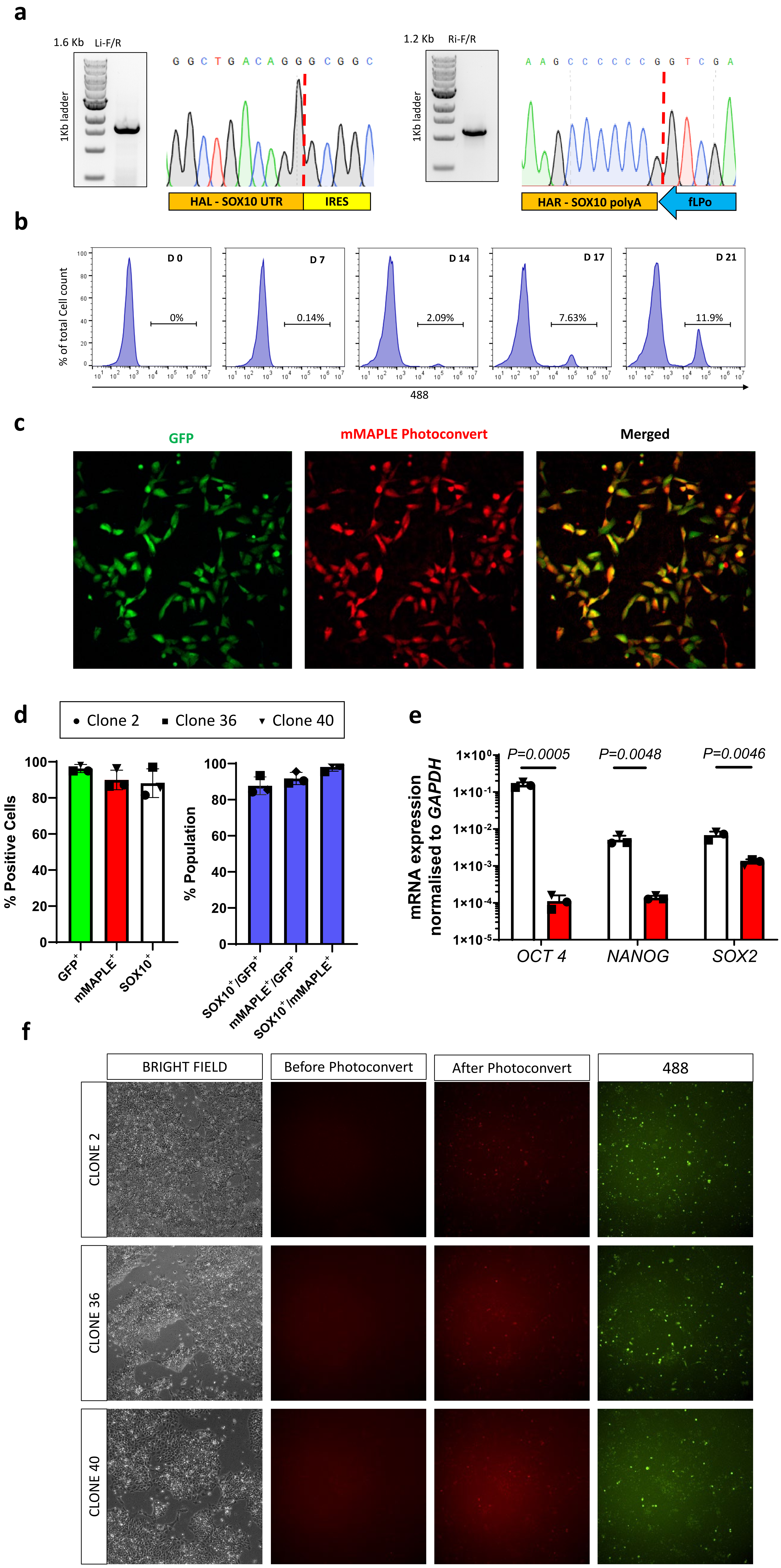

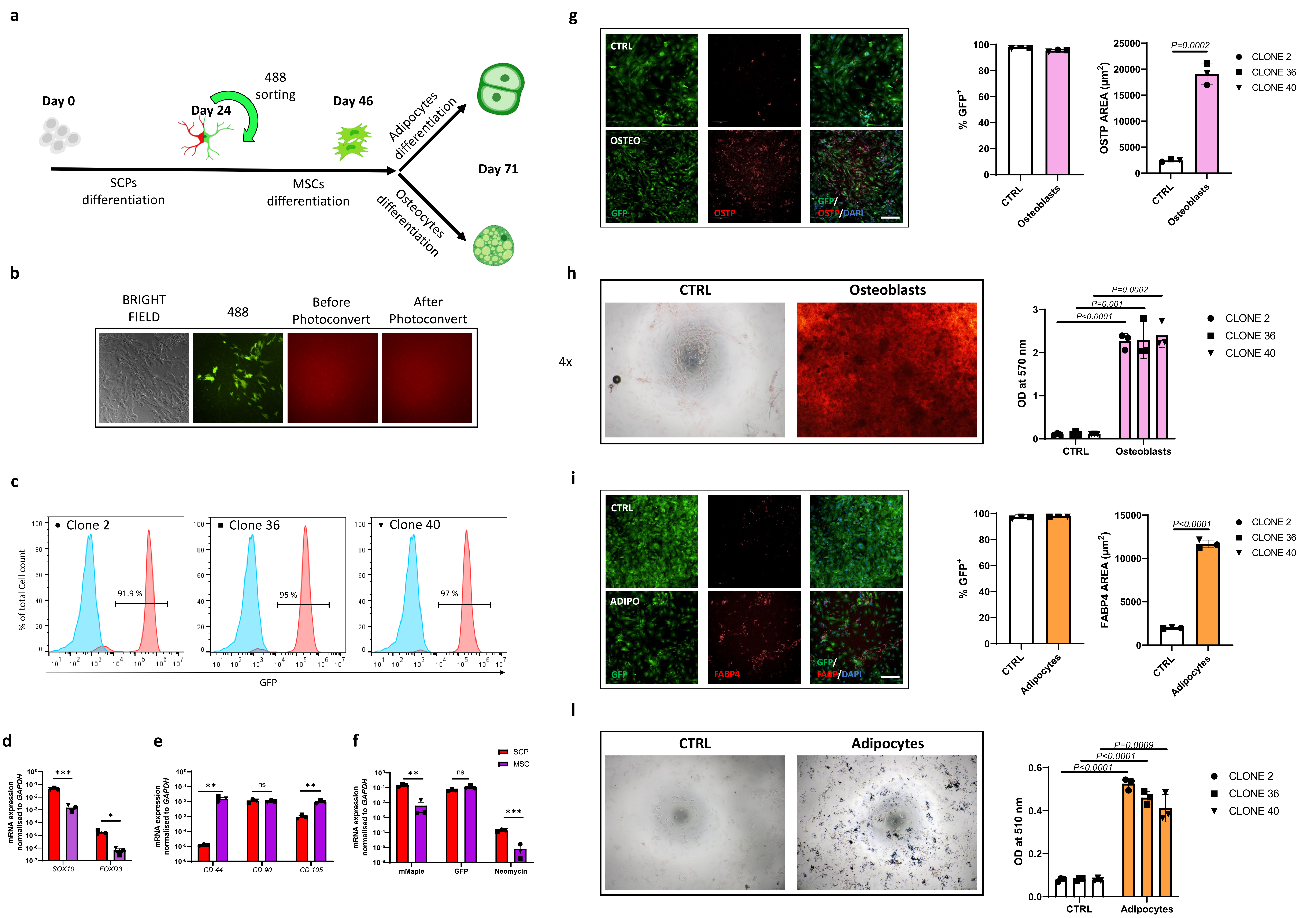

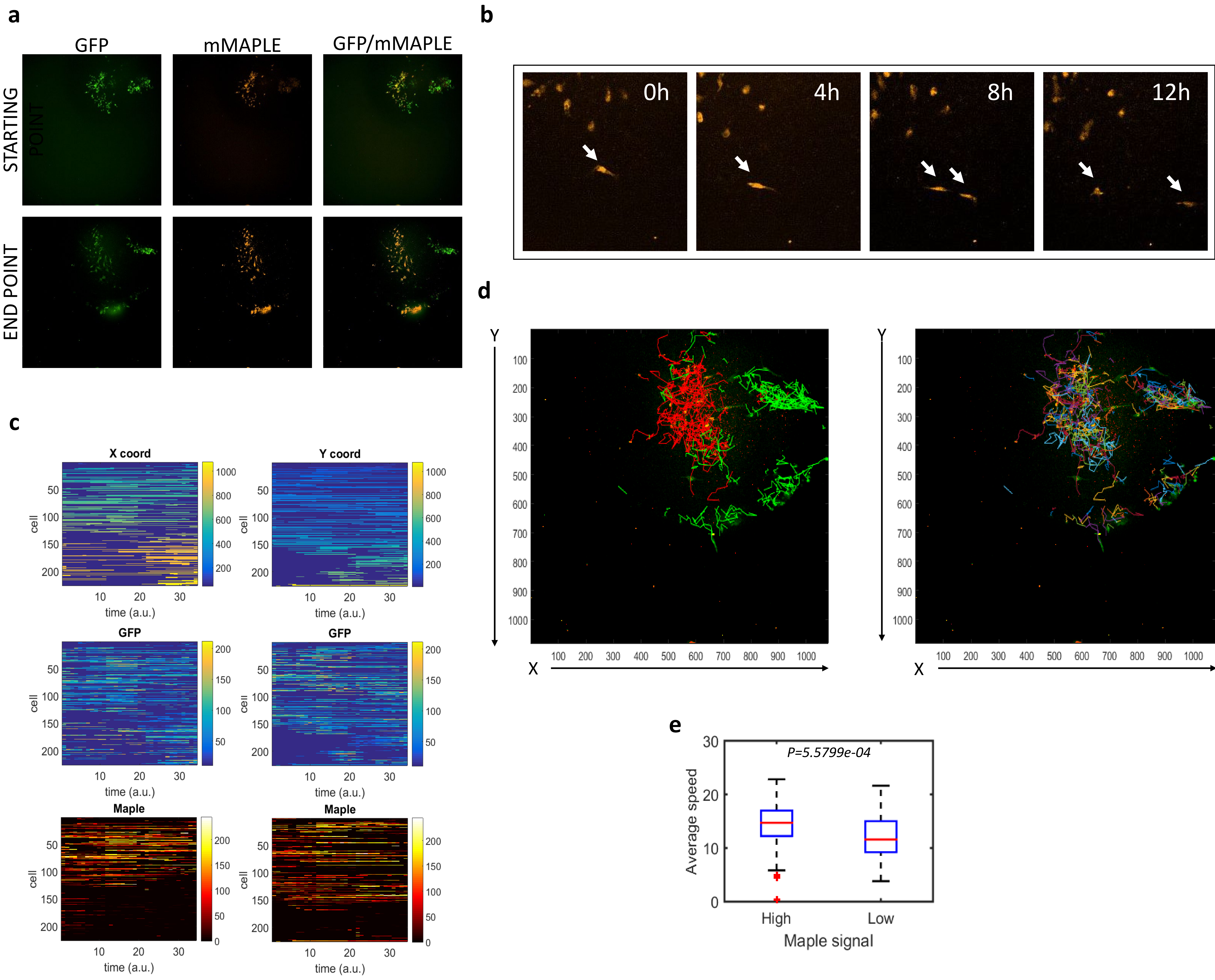

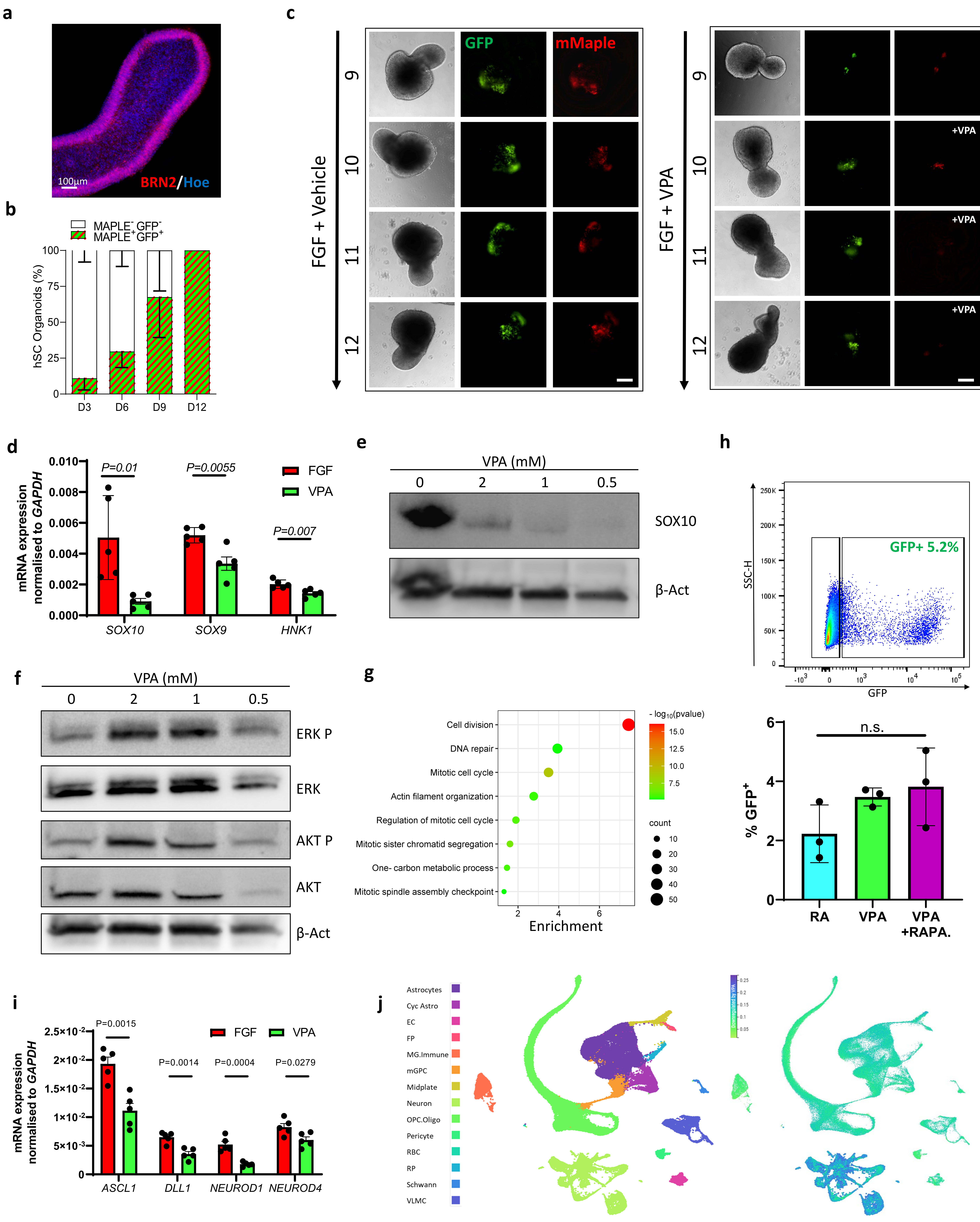

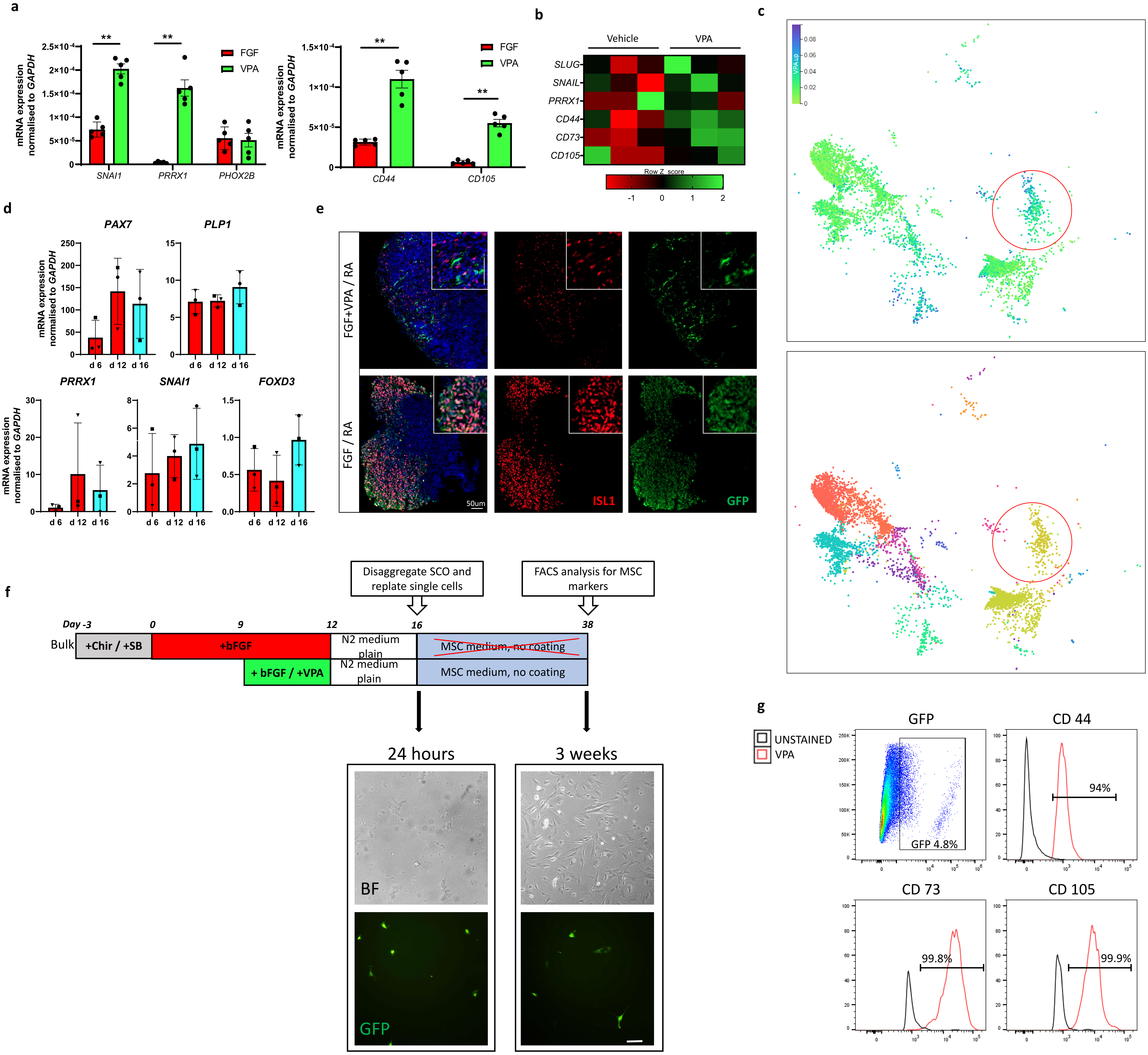

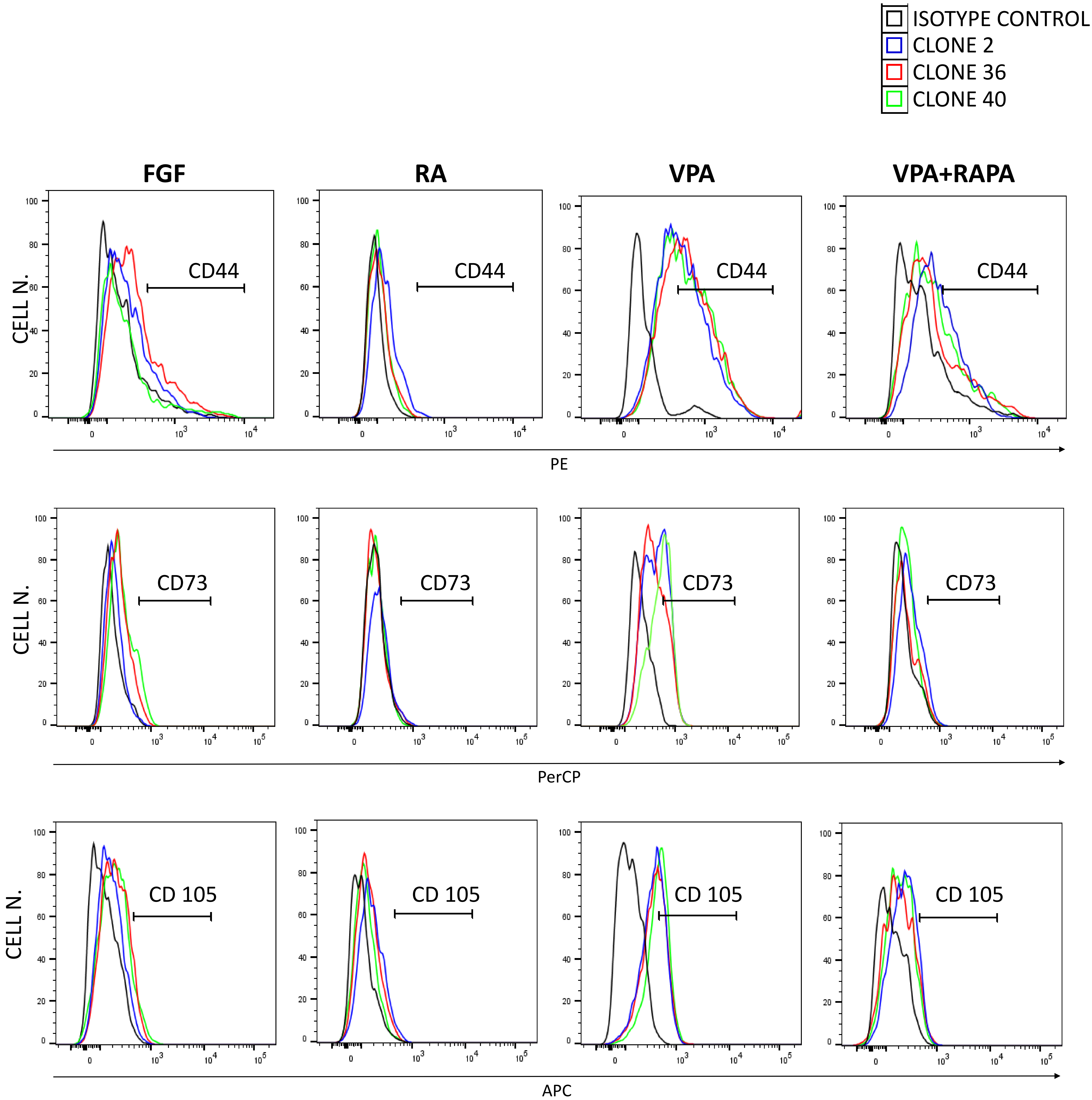

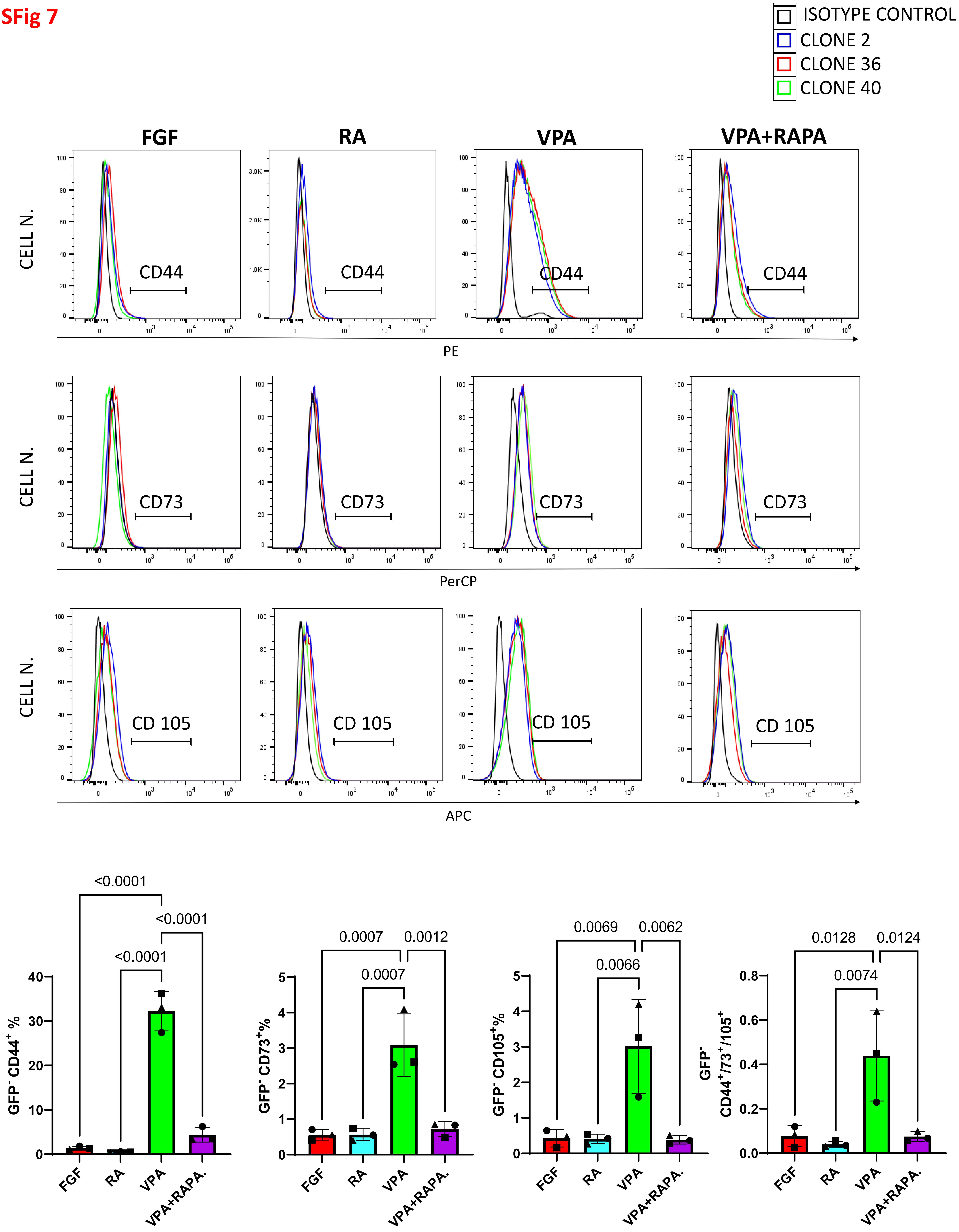

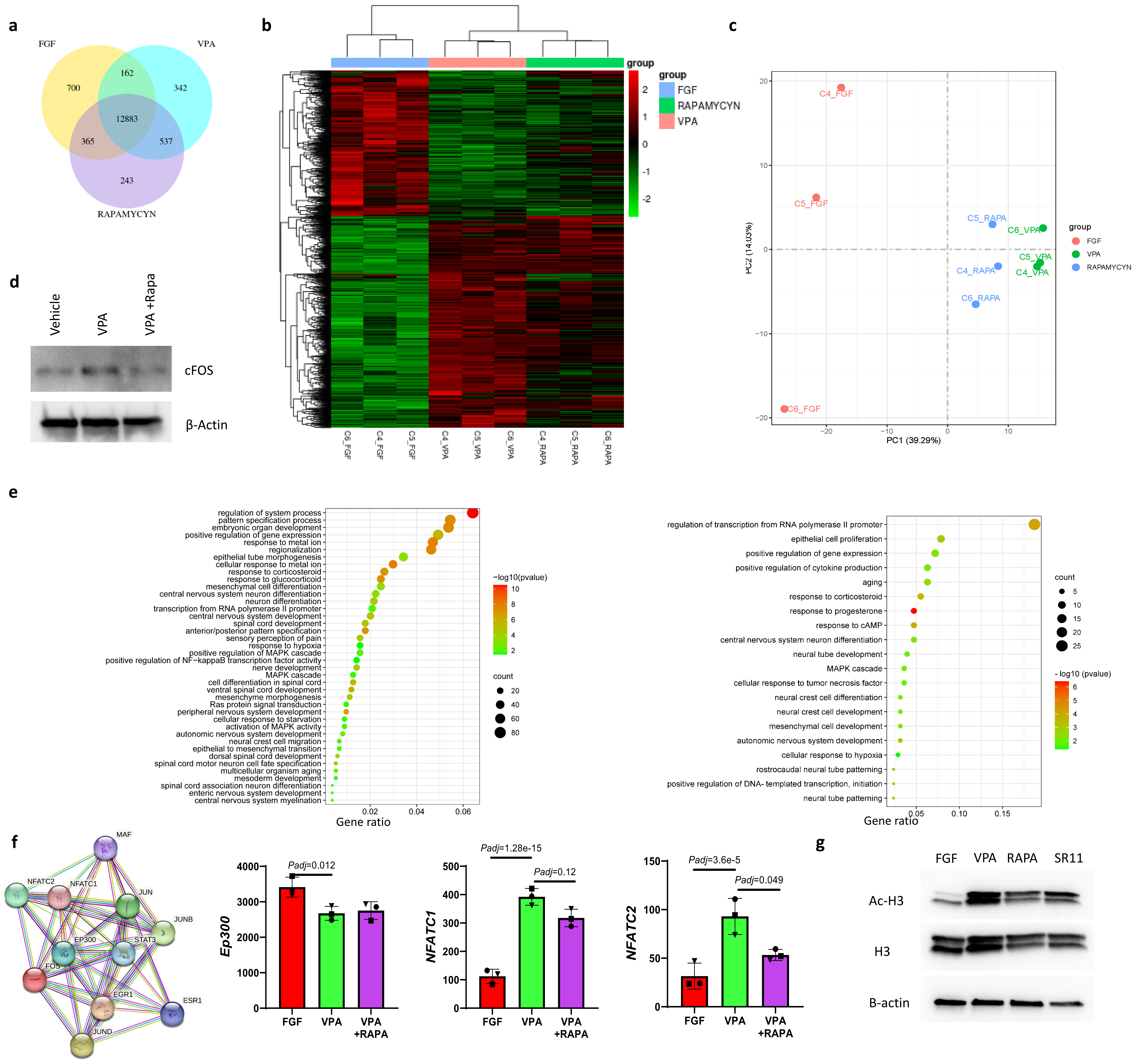
