## Supplementary material for "Rapamycin mitigates Valproic Acid-induced teratogenicity in human and animal models by suppressing AP-1-mediated senescence": Sup Table 1

| Primer | Sequence |
|---|---|
| Li F | CAGCAGAGAAGCTGATGAGAGGAAG |
| LiR | TAGGAATGCTCGTCAAGAAGACAGG |
| SeF | CAGATGCGACTTCAGAACCA |
| Se R | ATGTCGAAGCCTCAGGCTGTT |
| RiF | ACAAACGTGGTGGGCAACTGG |
| RiR | CTTTATTCAAATACCACTGGGAGGC |

| Gene | Direction | Sequence |
|---|---|---|
| *SOX10* | Forward | CATCCACCTCACAGATCGCC |
| | Reverse | GCATGTCAGACCCTCACTATCTG |
| *FOXD3* | Forward | GACGCAGGTTGCGATAGCC |
| | Reverse | CGCCTCCTTGGGCAATGTC |
| *CDH19* | Forward | ATCTGCACCCACTGGGACTT |
| | Reverse | CTGCTCAGGAACATGATGG |
| *MPZ* | Forward | GGTCCCCCACTTICTCAACC |
| | Reverse | TGTAAACCACGATGGCCTGG |
| *NGFR* | Forward | CCTACGGCTACTACCAGGATG |
| | Reverse | CACACGGTGTTCTGCTTGT |
| NEOMYCIN | Forward | AGACAATCGGCTGCTCTGAT |
| | Reverse | ATACTTTCTCGGCAGGAGCA |
| GFP | Forward | AAGGGCATCGACTTCAAGG |
| | Reverse | TGCTTGTCGGCCATGATATAG |
| mMaple | Forward | CAGATGCGACTTCAGAACCA |
| | Reverse | ATGTCGAAGCCTCAGGCTGTT |
| *SOX9* | Forward | AGGAAGTCGGTGAAGAACGG |
| | Reverse | CTGGGATTGCCCCGAGTG |
| *HNK1* | Forward | GCAGGTTGACGGCAAATCC |
| | Reverse | CCTGGCGTGGTCTACTTCG |
| *ASCL1* | Forward | CGCGGCCAACAAGAAGATG |
| | Reverse | CGACGAGTAGGATGAGACCG |
| *NEUROD1* | Forward | ATGACCAAATCGTACAGCGAG |
| | Reverse | GTTCATGGCTTCGAGGTCGT |
| *NEUROD4* | Forward | GAGAGCTAGTCAACACACCATC |
| | Reverse | GCATCCCATAAGTACCTGGTCTG |
| *DLL1* | Forward | GACGAACACTACTACGGAGAGG |
| | Reverse | AGCCAGGGTTGCACACTTT |
| *PHOX2B* | Forward | AACCCGATAAGGACCACTTTTG |
| | Reverse | AGAGTTTGTAAGGAACTGCGG |
| *HOXB1* | Forward | GAGCTTTGCACCGGCCTAT |
| | Reverse | CTTCATCCAGTCGAAGGTCCG |
| *OCT4* | Forward | GTACTCCTCGGTCCCTTTCC |
| | Reverse | CAAAAACCCTGGCACAAACT |
| *NANOG* | Forward | CAGCCCTGATTCTTCCACCAGTCCC |
| | Reverse | TGGAAGGTTCCCAGTCGGGTTCACC |

| | | |
|---|---|---|
| *SOX2* | Forward | TACAGCATGTCCTACTCGCAG |
| | Reverse | GAGGAAGAGGTAACCACAGGG |
| *CD44* | Forward | CTGCCGCTTTGCAGGTGTA |
| | Reverse | CATTGTGGGCAAGGTGCTATT |
| *CD90* | Forward | ATCGCTCTCCTGCTAACAGTC |
| | Reverse | CTCGTACTGGATGGGTGAACT |
| *CD105* | Forward | GCATCCTTCGTGGAGCTACC |
| | Reverse | GAGGAGTGGTCTGGATCGG |
| *SNAIL1* | Forward | TCGGAAGCCTAACTACAGCGA |
| | Reverse | AGATGAGCATTGGCAGCGAG |
| *PRRX1* | Forward | TGATGCTTTTGTGCGAGAAGA |
| | Reverse | AGGGAAGCGTTTTTATTGGCT |
| *PAX7* | Forward | GTCTCCAAGATTCTTTGCCG |
| | Reverse | CCACCTGTCTGGGCTTGCTG |
| *PLP1* | Forward | TGCTGATGCCAGAATGTATGG |
| | Reverse | GCAGATGGACAGAAGGTTGGA |
